# Remapping in cerebral and cerebellar cortices is not restricted by somatotopy

**DOI:** 10.1101/439356

**Authors:** Avital Hahamy, Tamar R. Makin

## Abstract

A fundamental organizing principle in the somatosensory and motor systems is somatotopy, where specific body parts are represented separately and adjacently to other body parts, resulting in a body map. Different terminals of the sensorimotor network show varied somatotopic layouts, in which the relative position, distance and overlap between body-part representations differ. Since somatotopy is best characterized in the primary somatosensory (S1) and motor (M1) cortices, these terminals have been the main focus of research on somatotopic remapping following loss of sensory input (e.g. arm amputation). Cortical remapping is generally considered to be driven by the layout of the underlying somatotopy, such that neighboring body-part representations tend to activate the deprived brain region. Here, we challenge the assumption that somatotopic layout restricts remapping, by comparing patterns of remapping in humans born without one hand (hereafter, one-handers, n=26) across multiple terminals of the sensorimotor pathway. We first report that in the cerebellum of one-handers, the deprived hand region represents multiple body parts. Importantly, the representations of some of these body parts do not neighbor the deprived hand region. We further replicate our previous finding, showing a similar pattern of remapping in the deprived hand region of the cerebral cortex in one-handers. Finally, we report preliminary results of a similar remapping pattern in the putamen of one-handers. Since these three sensorimotor terminals (cerebellum, cerebrum, putamen) contain different somatotopic layouts, the parallel remapping they undergo demonstrates that the mere spatial layout of body-part representations may not exclusively dictate remapping in the sensorimotor systems.

**Significance Statement:** When a hand is missing, the brain region that typically processes information from that hand may instead process information from other body-parts, a phenomenon termed remapping. It is commonly thought that only body-parts whose information is processed in regions neighboring the hand region could “take up” the resources of this now deprived region. Here we demonstrate that information from multiple body-parts is processed in the hand regions of both the cerebral cortex and cerebellum. The native brain regions of these body-parts have varying levels of overlap with the hand region across multiple terminals in the sensorimotor hierarchy, and do not necessarily neighbor the hand region. We therefore propose that proximity between brain regions does not limit brain remapping.

## Introduction

Somatotopic organization in primary somatosensory and motor cortices is thought to reflect the lateralized and segregated neural activation patterns associated with sensations from- and movements of-distinct body parts (Penfield and Boldrey, 1937; Penfield and Rasmussen, 1950; Catani, 2017; Roux et al., 2018). Following input and output loss, e.g. arm amputation in adults, S1/M1 somatotopies undergo remapping, such that the region previously representing the hand becomes responsive to inputs from other body-parts (Flor et al., 1995; Makin et al., 2013b; Chand and Jain, 2015; Raffin et al., 2016). The principles governing such architectural change are thought to derive from the underlying somatotopy: neighboring representations, which share greater cortical overlap (Merzenich et al., 1984; Pons et al., 1991; Merzenich and Jenkins, 1993; Florence et al., 1998) and/or receive stronger inhibition from now absent inputs (Faggin et al., 1997; Margolis et al., 2012) are more likely to activate the deprived cortical region. Subsequently, findings showing increased activation by facial inputs in the missing-hand region following arm amputation (interpreted as resulting from a presumed proximity between hand and lower-face representations (Jain et al., 2008; Kaas et al., 2008; MacIver et al., 2008; Foell et al., 2014; Andoh et al., 2017), have been taken as evidence for the role of somatotopy in scaffolding remapping.

We, and others, have recently challenged this view, by demonstrating that remapping may occur between both neighboring and distant body-part representations. For example, the intact hand of amputees shows increased activation in the S1/M1 missing-hand region (hereafter deprived cerebral hand region) (Bogdanov et al., 2012; Makin et al., 2013b; Philip and Frey, 2014). Similarly, individuals born without hands show increased feet activation in their deprived cerebral hand regions (Stoeckel et al., 2009; Yu et al., 2014; Striem-Amit et al., 2018). This feet-to-hands remapping occurs despite the inherent cortical distance between the native regions of the feet and hands. Finally, in individuals born without one hand (hereafter, one-handers), multiple body-parts (residual arm, lips and feet, but not the intact hand) activate the deprived cerebral hand region (Makin et al., 2013b; Hahamy et al., 2017). As the native foot and lip regions are not immediately neighboring the hand region, we suggested that proximity between body-part representations is not a prerequisite for remapping. Yet, it has recently been argued that in cases of congenital hand-loss, remapping is driven by topographic constraints, such that body-part representations that are further from the hand region will show reduced remapping compared to representations that neighbor the hand region (Striem-Amit et al., 2018). Thus, local somatotopy is still considered the main driver of remapping in both congenital and late-onset sensorimotor deprivation.

Here, we address this question by examining remapping in one-handers, by measuring the level of activation in the deprived hand region evoked by movements of multiple body-parts. Crucially, remapping is examined at multiple sensorimotor terminals with varying somatotopies. Somatotopic organization was previously identified throughout the sensorimotor systems, including the cerebellum (Manni and Petrosini, 2004), brainstem (Jang et al., 2011) and basal ganglia (Nambu, 2011). Here, we focus on the cerebellum, where somatotopy can be reliably identified using fMRI (Grodd et al., 2001; Buckner et al., 2011; Wiestler et al., 2011; Haak et al., 2017). The cerebellum’s somatotopy differs from that of S1/M1. For example, in S1/M1 the hand and arm have separate regions, and the lip and foot regions are equally distant from the hand region (Makin et al., 2015). However, in the cerebellum, the arm and hand regions are overlapping, and the lip region partially overlaps with the hand region (Manni and Petrosini, 2004; Mottolese et al., 2013; Mottolese et al., 2015). If mere somatotopy drives remapping (Flor et al., 1995; Chand and Jain, 2015; Raffin et al., 2016; Striem-Amit et al., 2018), then these different somatotopies should result in different remapping patterns between the cerebral and cerebellar deprived hand regions. However, if similar patterns of remapping would be observed across these terminals, it is less likely that remapping is solely determined by the local somatotopy (Hahamy et al., 2017). We test these competing hypotheses using several independently-acquired datasets of one-handers and two-handed controls and a meta-analysis approach. Our findings provide robust evidence for similar body-part remapping across the hierarchy of the sensorimotor system. We therefore propose that remapping is not necessarily restricted by the physical proximity between the native and remapped representations, and discuss alternative factors that may underlie this extensive brain plasticity.

## Materials and Methods

To avoid known issues of flexibility in fMRI analyses (Carp, 2012) and to enable replication, we harmonized our methods, including experimental design, preprocessing steps and statistical analyses across datasets to compare with our previous publication (Hahamy et al., 2017).

### Participants

This study makes use of three independently acquired fMRI datasets, each containing data of both one-handers and two-handed controls. Two of these datasets had cerebellar coverage, and were therefore used for cerebellar analyses. All three datasets were used for analysis of the cerebral cortex (cerebral cortex findings in the third dataset have been published in Hahamy et al., 2017).

Recruitment was carried out in accordance with NHS national research ethics service approval (10/H0707/29, dataset1) and with Oxford University’s Medical Sciences inter-divisional research ethics committee (MS-IDREC-C2-2015-012, dataset2). Informed consent and consent to publish was obtained in accordance with ethical standards set out by the Declaration of Helsinki.

The first dataset (hereafter, Dataset1) contained the same population recruited for a previous study, using the same scanning procedures and exclusion criteria as described before (Hahamy et al., 2015b). 25 healthy controls (15 females, age = 41.12±12.86, 8 left hand dominant) and 14 individuals with a congenital unilateral upper limb deficit (one-handers, 9 females, age = 36.64±12.02, 4 with absent right hand) were recruited for the study. The proportion of one-handers with a missing right hand (n=4) and controls who are left-hand dominant (n=8) were similar (χ^2^_(1)_ =0.18, p=0.67).

The second dataset (hereafter, Dataset2) was acquired as part of a larger study (the full study protocol is currently under preparation and will be made available via Open Science Framework). These data included the scanning of 12 healthy controls (5 females, age = 45.33±14.85, 5 left hand dominant), and 14 one-handers (7 females, age = 45.25±11.38, 6 with absent right hand) (see Table 1 for demographic details). The proportions of one-handers with a missing right hand (n=6) and controls who are left-hand dominant (n=5) were similar (χ^2^_(1)_ =0.05, p=0.82). Four one-handers participated in both studies, with data acquired approximately 5 years apart.

**Table 1.**
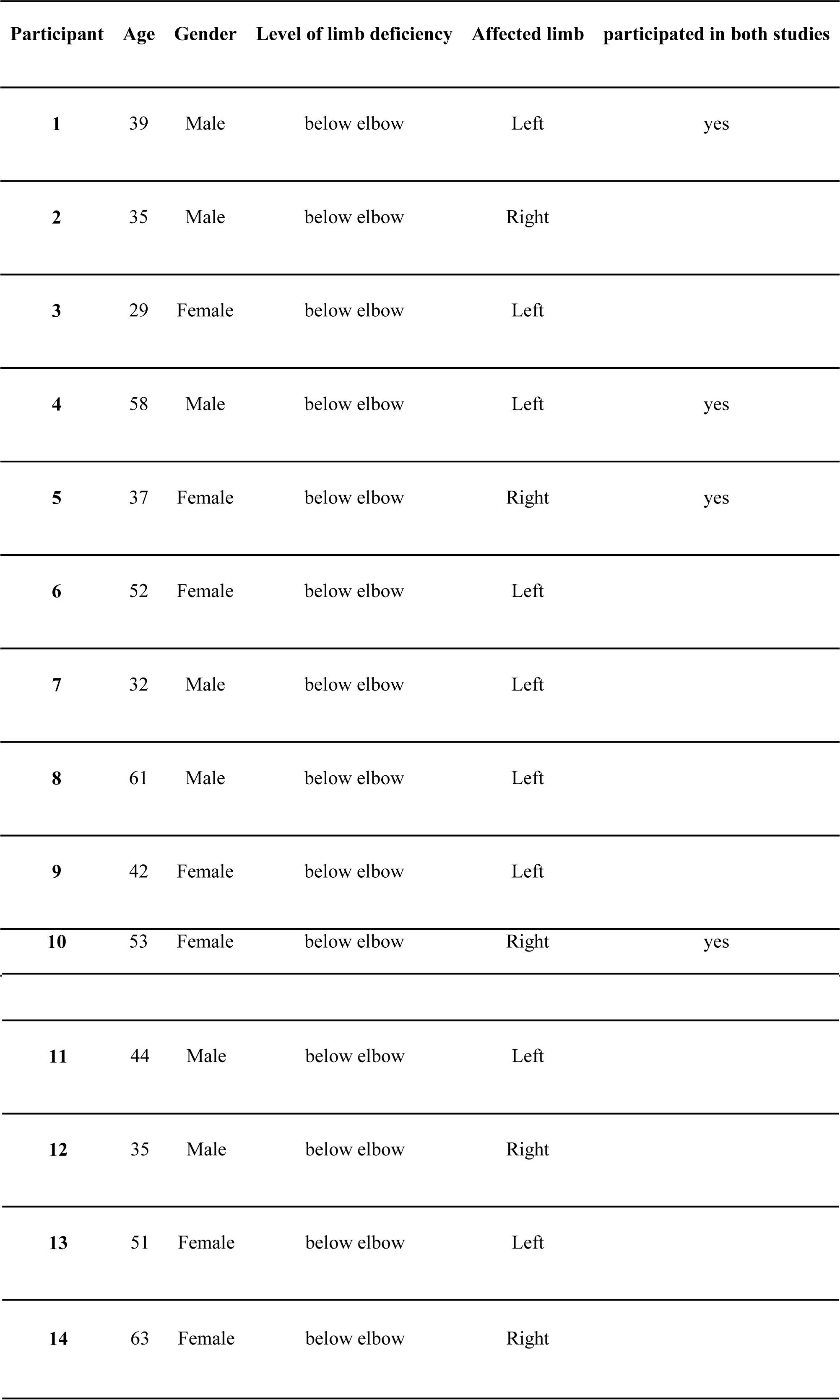
Demographic details of individuals with congenital limb-absence included in Dataset2 (Full details of the participants of Dataset1 are available in Hahamy et al., 2015b).

| Participant | Age | Gender | Level of limb deficiency | Affected limb | participated in both studies |
| --- | --- | --- | --- | --- | --- |
| 1 | 39 | Male | below elbow | Left | yes |
| 2 | 35 | Male | below elbow | Right |  |
| 3 | 29 | Female | below elbow | Left |  |
| 4 | 58 | Male | below elbow | Left | yes |
| 5 | 37 | Female | below elbow | Right | yes |
| 6 | 52 | Female | below elbow | Left |  |
| 7 | 32 | Male | below elbow | Left |  |
| 8 | 61 | Male | below elbow | Left |  |
| 9 | 42 | Female | below elbow | Left |  |
| 10 | 53 | Female | below elbow | Right | yes |

|  |  |  |  |  |
| --- | --- | --- | --- | --- |
| **11** | 44 | Male | below elbow | Left |

|  |  |  |  |  |
| --- | --- | --- | --- | --- |
| **12** | 35 | Male | below elbow | Right |

|  |  |  |  |  |
| --- | --- | --- | --- | --- |
| **13** | 51 | Female | below elbow | Left |

|  |  |  |  |  |
| --- | --- | --- | --- | --- |
| **14** | 63 | Female | below elbow | Right |

Full demographic description and acquisition-related information regarding the third dataset are available (Hahamy et al., 2017).

### Experimental Design

Scanning protocol for both datasets included multiple scans (see protocol in https://osf.io/4vcmx/). Only an anatomical T1 scan and a task scan for body-part functional localization were used and analyzed here (these scanning procedures were described previously, see Makin et al., 2013b; Hahamy et al., 2015b).

The sensorimotor task in both datasets followed the same procedure: Participants were visually instructed to move each of their hands (finger flexion/extension), arms (elbow flexion/extension), their feet (bilateral toe movements), or lips, as paced by a visual cue. None of the one-handers experienced phantom sensations. Therefore, in conditions concerning missing hand movements (and elbow movements for one participant with an above-elbow deficiency) participants were instructed to imagine moving their missing limb. This condition was only included to match the experimental design across groups and was not used for main analysis. The protocol consisted of alternating 12-s periods of movement and rest. Each of the six conditions was repeated four times in a semi-counterbalanced order. Participants were trained before the scan on the degree and form of the movements. To confirm that appropriate movements were made at the instructed times, task performance was visually monitored online, and video recordings were made in a subset of the scans for further off-line evaluation.

### MRI data acquisition

The MRI measurements of Dataset1 were obtained using a 3T Verio scanner (Siemens, Erlangen, Germany) with a 32-channel head coil. Anatomical data were acquired using a T1-weighted magnetization prepared rapid acquisition gradient echo sequence (MPRAGE) with the parameters: TR: 2040 ms; TE: 4.7 ms; flip angle: 8°, voxel size: 1 mm isotropic resolution. Functional data based on the blood oxygenation level-dependent (BOLD) signal were acquired using a multiple gradient echo-planar T2*-weighted pulse sequence, with the parameters: TR: 2000 ms; TE: 30 ms; flip angle: 90°; imaging matrix: 64 × 64; FOV: 192 mm axial slices. 46 slices with slice thickness of 3 mm and no gap were oriented in the oblique axial plane, covering the whole cortex, with partial coverage of the cerebellum.

MRI images of Dataset2 were acquired using a 3T MAGNETON Prisma MRI scanner (Siemens, Erlangen, Germany) with a 32-channel head coil. Anatomical images were acquired using a T1-weighted sequence with the parameters TR: 1900 ms, TE: 3.97 ms, flip angle: 8°, voxel size: 1 mm isotropic resolution. Functional images were collected using a multiband T2*-weighted pulse sequence with a between-slice acceleration factor of 4 and no in-slice acceleration. This allowed acquiring data with increased spatial (2mm isotropic) and temporal (TR: 1500ms) resolution, covering the entire brain. The following acquisition parameters were used - TE: 32.40ms; flip angle: 75°, 72 transversal slices. Field maps were acquired for field unwarping.

### Preprocessing of functional data

All imaging data were processed using FSL 5.1 (www.fmrib.ox.ac.uk/fsl). Data collected for individuals with absent right limbs were mirror reversed across the mid-sagittal plane prior to all analyses so that the hemisphere corresponding to the missing hand was consistently aligned. Data collected for left-hand dominant controls were also flipped, in order to account for potential biases stemming from this procedure. The proportion of flipped data did not differ between experimental groups in either dataset (χ^2^_(1)_) =0.18, p=0.67 for Dataset1; χ^2^_(1)_ =0.05, p=0.82 for Dataset2), and this flipping procedure has been validated using multiple approaches (see Hahamy et al., 2017).

Functional data were analyzed using FMRIB’s expert analysis tool (FEAT, version 5.98). The following pre-statistics processing was applied to each individual task-based run: motion correction using FMRIB’s Linear Image Registration Tool (Jenkinson et al., 2002); brain-extraction using BET (Smith, 2002); mean-based intensity normalization; high pass temporal filtering of 100 s; and spatial smoothing using a Gaussian kernel of FWHM (full width at half maximum) 4 mm. Time-course statistical analysis was carried out using FILM (FMRIB’s Improved Linear Model) with local autocorrelation correction. Functional data were aligned to structural images (within-subject) initially using linear registration (FMRIB’s Linear Image Registration Tool, FLIRT), then optimized using Boundary-Based Registration (Greve and Fischl, 2009). Structural images were transformed to standard MNI space using a non-linear registration tool (FNIRT), and the resulting warp fields were applied to the functional statistical summary images.

### Statistical analyses

#### Meta-analysis approach

The current study makes use of three separate datasets, acquired across several years and using different magnets and scanning parameters. Two of these datasets included coverage of the cerebellum and were therefore used for cerebellar analysis, and all three datasets were used for analysis of the cerebral cortex. Multiple datasets can, in principle, be collapsed for analysis purposes, benefiting from statistical power to identify weak effects that may not be noticeable in each separate dataset (Friston, 2012). However, as the current study is guided by an a-priori hypothesis that is also spatially focal (remapping in the deprived hand region of one-handers), it calls for more stringent inference methods rather than for exploratory ones that benefit from enhanced power. We therefore opted to analyzing each dataset separately and combine results using a meta-analysis approach (Hahamy et al., 2015a). Differences across datasets are naturally expected, given the inherent variability between datasets (different scanners/scanning protocols/participants) as well as various noise factors that influence any fMRI measurement. However, while inter-dataset variability could be attributed to both noise and experiment-related phenomena, consistent effects across datasets can only be attributed to the latter. Thus, the use of meta-analysis allowed us to test the inherent reproducibility of findings across datasets, and hence make more valid inferences (Ioannidis et al., 2014; Picciotto, 2018).

Our analysis pipeline for all ROI-based analyses reported below included a-parametric permutation tests performed within each separate dataset. Permutation tests are statistically stringent as they make no assumptions regarding the finite sample distribution of the data, but rather derive it given the data observed (Holmes et al., 1996; Nichols and Holmes, 2002), and are also less sensitive to outlier effects (Masyn et al., 2013), thus contributing to the robustness of findings. The dataset–specific p-values resulting from each of the below-described permutation tests were then combined across datasets and meta-analyzed using Fisher’s method (Fisher, 1925; Fisher, 1948) to test the reproducibility of results across datasets. In order to establish the robustness of the reported effects, p-values were additionally tested using Stouffer’s test (Stouffer et al., 1949) and the weighted Z-test (weights set to the square root of each sample size, Liptak, 1958). To correct for multiple hypotheses testing across the 3 experimental conditions of interest (movements of the residual arm, lips and feet), the alpha level was adjusted to 0.017 based on the highly conservative Bonferroni correction.

#### Whole-brain analysis

To evaluate whether movements of different body-parts differentially activate the brains of one-handers compared to controls, activation evoked by movements of these different body-parts was compared between the experimental groups. Movements of the lips and feet were directly compared between groups, intact hand movements in the one-handed group were compared with dominant hand movements in controls, and residual arm movements in the one-handed group were compared with non-dominant arm movements in controls. All statistical analyses were designed to follow the procedures described in our original report (Hahamy et al., 2017). Statistical analyses were conducted using FSL and in-house Matlab code. To compute task-based statistical parametric maps, we applied a voxel-based general linear model (GLM), as implemented in FEAT, using a double-gamma hemodynamic response function and its temporal derivative convolved with the experimental model. The 6 motion parameters and their derivatives were also included in the GLM as nuisance regressors. Our main comparisons contrasted intact/dominant hand, residual/nondominant arm, lips and feet conditions against a baseline (rest) condition.

Second-level analysis of statistical maps was carried out using FMRIB’s Local Analysis of Mixed Effects (FLAME). The cross-subject GLM included planned comparisons between the two groups. Z (Gaussianized T/F) statistic images were thresholded using clusters determined by Z>2.6 (p<0.01), and a family-wise-error corrected cluster significance threshold of p<0.01 was applied to the suprathreshold clusters. This whole-brain analysis tests the specificity of plasticity to the deprived hand region of one-handers, hence a lenient statistical threshold (p<0.05) is typically used in such procedures (Makin et al., 2013b; Hahamy et al., 2017). Nevertheless, as we test several whole-brain comparisons (residual arm, lips and feet conditions), we chose a more strict threshold of 0.01 across our tests to correct for any alpha inflation. The nature of the sensorimotor task, in combination with the spatial acquisition resolution, the smoothing and coregistration steps, precludes us from reliably separating sensory and motor sub-regions. As such, all results are regarded as ‘sensorimotor’.

For visualization purposes only, condition-specific within-group maps were created for both the cerebellum and cerebral cortex, using the same statistical procedures reported above. These maps were merely aimed at visualizing the sources of the reported group-differences, and hence were presented at varying thresholds that best capture the effects observed in the direct statistical comparisons between groups. Specifically, all maps were thresholded at p<0.01, except for the cerebral maps of the arm condition, which were thresholded at of p<0.0006. Since arm movements massively activate the hand region, the choice of a more stringent threshold for these maps enabled a better visualization of the group-differences in overlap between peak activity and the deprived hand region. Using the same rationale, maps were presented before correction for multiple comparison, to best visualize group differences.

For presentation purposes, statistical parametric activation maps of the cerebellum were projected onto a flat cerebellar surface using SUIT (Diedrichsen and Zotow, 2015), and parametric activations in the cerebral cortex were projected onto an inflated cortical surface of a representative participant’s cortex using the Connectome Workbench.

#### Cerebellar regions of interest (ROI) definition

To ascertain that the observed increased cerebellar activation in one-handers (observed across the two datasets with cerebellar coverage) falls within the hand region, and to measure its extent, single-subject activation values were extracted from independently defined hand-region ROIs and compared between experimental groups. Activations in the control group of each dataset were used to define ROIs for the second dataset (thus keeping the ROI definition independent of the tested data). Thus, to define the cerebellar hand regions of the dominant/intact hemisphere (ipsilateral to the dominant/intact hand) and nondominant/deprived hemisphere (contralateral to the dominant/intact hand), the 100 cerebellar voxels of highest activation evoked by either dominant or nondominant hand movements in the control group of one dataset were used as ROIs in the second dataset, and vice-versa (see Table 2, Figures 1,2). Percent signal change activation values from the individual statistical parametric maps were extracted for the intact and deprived hand ROIs for each participant in the residual/nondominant arm, lips, feet and intact hand conditions. Since the functional data of one control participant in Dataset1 did not cover the cerebellum, data from this individual were excluded from the cerebellar ROI analysis. The same method was used to define the cerebellar regions of the lips and feet for visualization purposes.

**Table 2.**
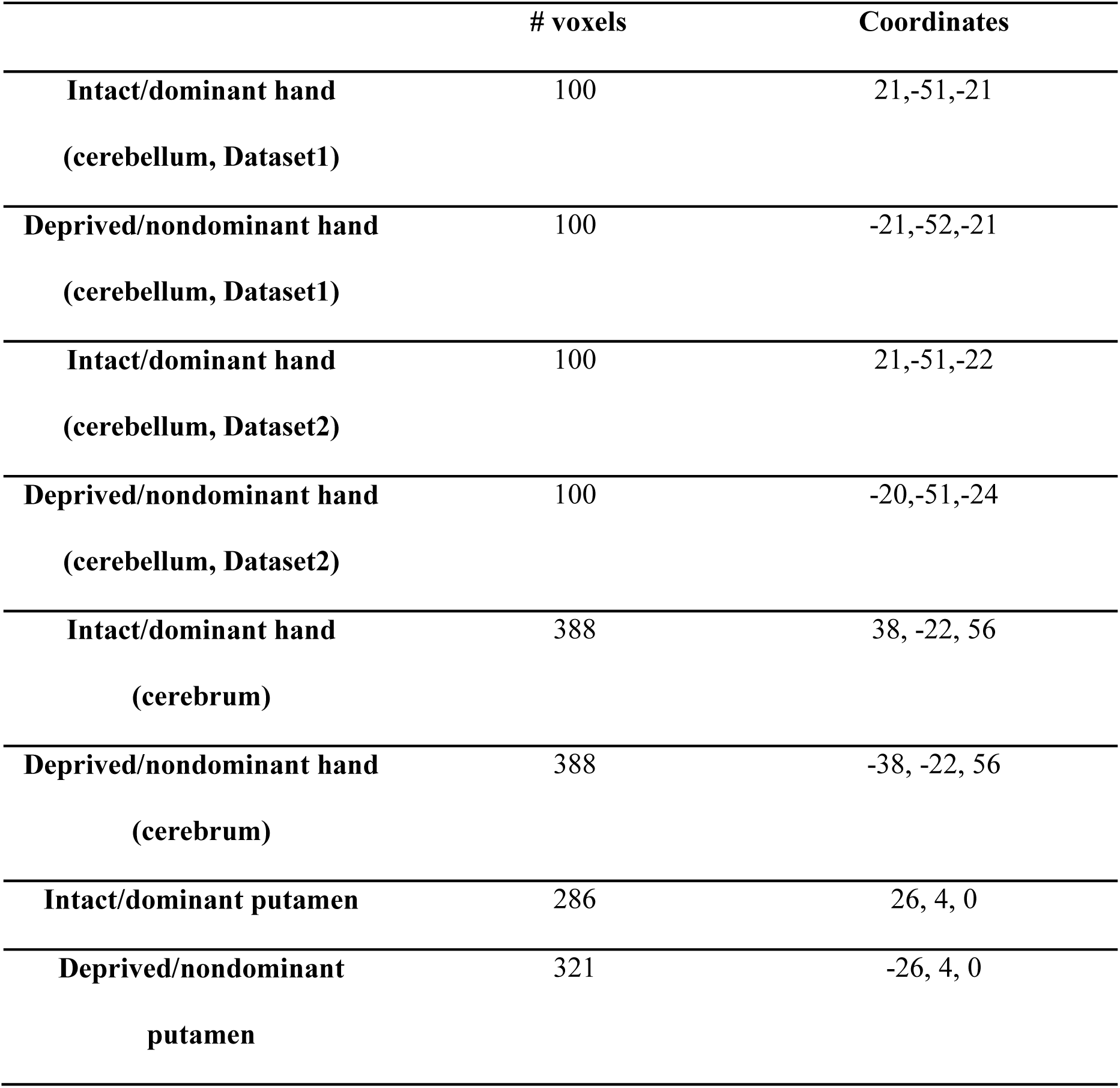
Number of voxels (2mm^3^) and center-of-gravity coordinates of regions of interest.

|  | # voxels | Coordinates |
| --- | --- | --- |
| **Intact/dominant hand**<br>**(cerebellum, Dataset1)** | 100 | 21,-51,-21 |
| **Deprived/nondominant hand**<br>**(cerebellum, Dataset1)** | 100 | -21,-52,-21 |
| **Intact/dominant hand**<br>**(cerebellum, Dataset2)** | 100 | 21,-51,-22 |
| **Deprived/nondominant hand**<br>**(cerebellum, Dataset2)** | 100 | -20,-51,-24 |
| **Intact/dominant hand**<br>**(cerebrum)** | 388 | 38, -22, 56 |
| **Deprived/nondominant hand**<br>**(cerebrum)** | 388 | -38, -22, 56 |
| **Intact/dominant putamen** | 286 | 26, 4, 0 |
| **Deprived/nondominant**<br>**putamen** | 321 | -26, 4, 0 |

**Figure 1.**
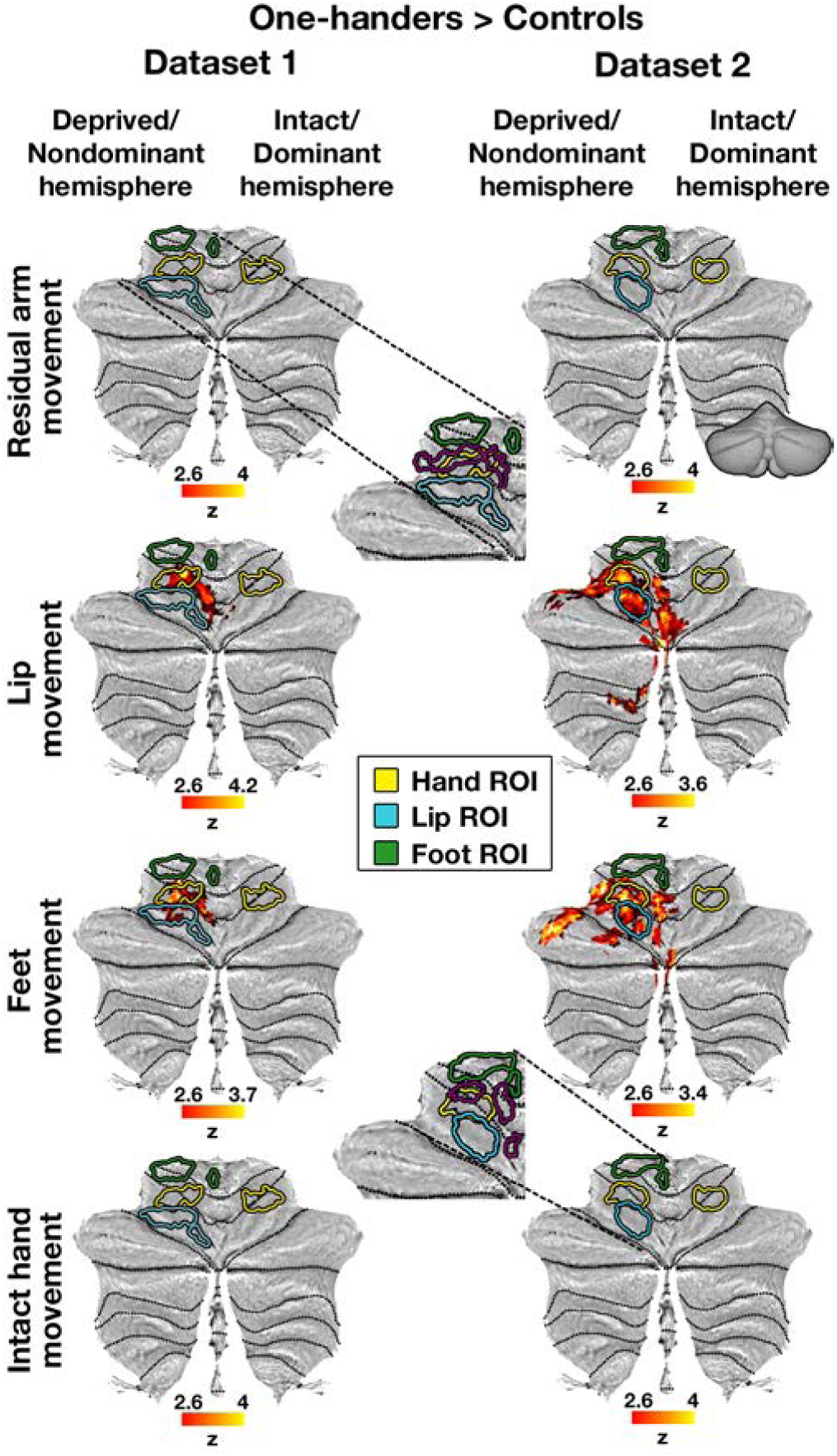
Representation of multiple body-parts in the deprived cerebellar hand region of one-handers: between-group contrast maps. The left/right panels show between-group contrast maps of Dataset1/Dataset2, respectively, during residual/nondominant arm (one-handers/controls), lips, feet and intact/dominant hand movements, projected onto a flat surface of the cerebellum (see example of an inflated surface on the top right). In the lips and feet conditions (but not in the residual arm or intact hand conditions), one-handers showed increased activation compared to controls, centred on the deprived cerebellar hand region. Yellow/blue/green contours indicate the hand/lip/foot ROIs, respectively. Inserts in the middle panel show the independent ROIs used in each of the datasets, defined based on the activations of the other dataset to ensure statistical independence. Middle inserts also include purple contours indicating the residual arm region. Intact/dominant hemisphere, ipsilateral to the intact/dominant hand; deprived/nondominant hemisphere, ipsilateral to missing/nondominant hand. All maps were cluster-based corrected for multiple comparisons across the entire brain. Results of residual arm movements in a subset of participants from Dataset1 were previously reported (Makin et al., 2013b).

**Figure 2.**
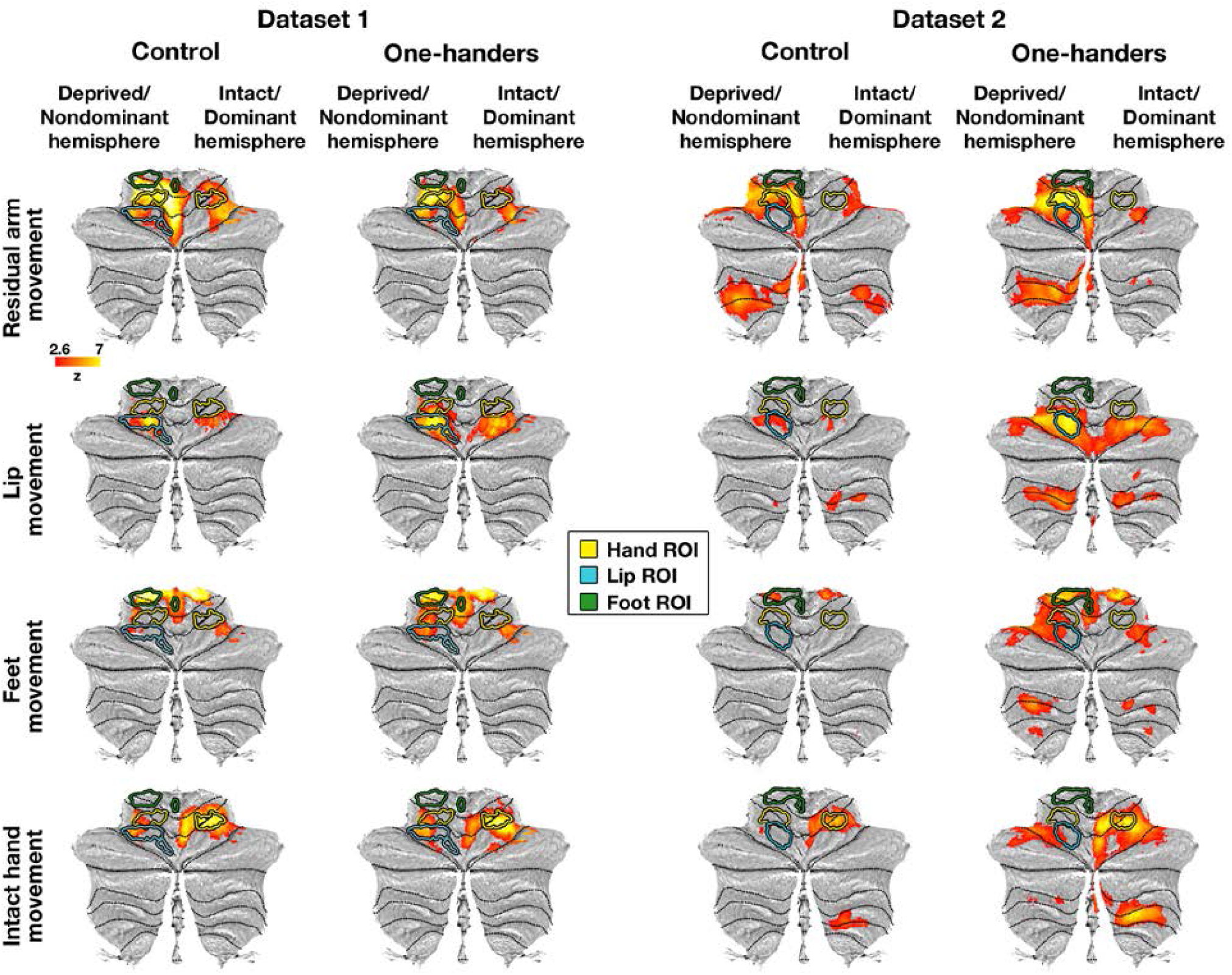
Cerebellar within-group activation maps. Within-group activation maps for each experimental condition versus a resting baseline (rows) are presented for the control and one-handed groups of each separate dataset (columns). All annotations are as detailed in Figure 1. All maps are presented at an uncorrected threshold of p<0.01 to visualize the origin of the between-group contrast results presented in Figure 1. The within-group maps of one-handers show over-activation in the cerebellar deprived hand region in the residual arm, lips and feet conditions, compared to controls.

#### Statistical analysis of cerebellar ROIs

To a-parametrically assess each planned group-contrast (experimental conditions involving movements of different body parts), permutation tests were employed within each dataset separately (Holmes et al., 1996; Nichols and Holmes, 2002). In each experimental condition separately, the test statistic was set as the difference between mean group activations in a certain ROI. Next, participants’ labels (one-handers or controls) were permuted under the null hypothesis of no group-differences in the levels of ROI activation under each experimental condition. Thus, two random experimental groups were created for each condition, and the difference between the groups’ mean activation in a given ROI was calculated. This procedure was repeated 10,000 times, creating 10,000 random differences that constructed the null distribution. For each experimental condition, the position of the true (unshuffled) group-difference relative to the null distribution was used to obtain a two-sided p-value. Using the same pipeline, under the null hypothesis of no 2-way interactions between groups and hemispheres (ipsilateral and contralateral to the missing/nondominant hand), both participants’ labels and within-participant hemisphere-labels were permuted in each dataset and experimental condition separately. The differences between hemisphere-scores were calculated per participant, averaged across participants of the same experimental group, and mean group differences were derived. The position of the true (unshuffled) group-difference relative to the null distribution (resulting from 10,000 such iterations) in each experimental condition was used to derive a two-sided p-value. Note that due to the somatotopic arrangement within the sensorimotor terminals, in which the hand and arm representations overlap, movements of the hand and arm evoke much higher activation in the hand region compared to movements of the lips and feet in the typical brain. These known differences in activation levels in the hand-region preclude us from running a formal direct comparison across body-parts, such as a 3-way ANOVA.

To comparatively examine the level of remapping of the foot and lip representations (which do not natively overlap with the hand representation), the activations evoked by lips and feet movements in the deprived cerebellar hand region of one-handers were directly compared using a permutation test, following the same procedure previously detailed. Furthermore, a 3-way interaction with factors group (controls/one-handers), hemisphere (intact/deprived) and body-part (lips/feet) was calculated.

#### Assessing remapping in the cerebral cortex

The overlap of participants between Dataset1 and Dataset2 is relatively small (4 out of 24 participants), which allowed us to perform the above described cross-dataset replication analyses for the cerebellum. However, we also aimed to test the reproducibility of our previously reported findings of remapping in the cerebral cortex (Hahamy et al., 2017) using the two current datasets and a previously published dataset, and these contained a larger overlap of participants. Dataset2 included only 5 participants who also participated in the Hahamy et al., 2017 study (with data acquired approximately 2 years apart). However, Dataset1 and the data used in Hahamy et al., 2017 greatly overlapped (12 out of 14 participants, with data acquired approximately 3 years apart). Hence, cerebral-related results obtained from Dataset1 should be taken as a measure of a within-group replication over time with regards to Hahamy et al., 2017, rather than as a between-group replication over participants.

Analyses performed on the cerebral cortex were identical to those described for the cerebellum, except for the following differences: 1) For cerebral hand ROIs (in both hemispheres) we used independent regions previously defined based on the original sample of one-handers and controls of our previous study (Hahamy et al., 2017; see Table 2; these ROIs will be made freely available via open science framework), to standardize the analysis across the three datasets (the two current ones and the previously published one). ROIs for lips and feet were also adopted from the same previous work to visualize the S1/M1 somatotopy. We were unable to reliably separate between the two cerebral foot regions for two reasons. First, our experimental task comprised of simultaneous movements of the two feet. Second, the resulting activation in the two foot regions occupied the medial surface of the cerebral cortex, mixing signals from the two hemispheres due to acquisition, preprocessing and coregistration parameters. For this reason, a bilateral ROI was defined for the feet. 2) Since the cerebral-focused ROI analyses were guided by a predefined hypothesis (over-representation of the residual arm, lips and feet in the deprived hand region of one-handers) based on our previous study, one-tailed statistical tests within each of Dataset1 and Dataset2 were used. Since our previous study did not find a significant interaction between groups and hemispheres for the feet condition, the tests of this effect in the two current datasets were performed in a two-tailed form.

For each experimental condition, sets of 3 dataset-specific p-values (resulting from each of the two current datasets, as well as the previously reported p-values of the original dataset) were combined and tested using the same methods described for the cerebellum analyses, including correction for multiple comparisons.

#### Comparing remapping in the cerebellar and cerebral cortices

To compare the relative levels of remapping of the lips and feet into the cerebrum and cerebellum hand regions, the ratio between lips-evoked and feet-evoked activation was calculated in the deprived hand region of the cerebrum and cerebellum for each one-hander. The cerebrum remapping ratio was then divided by the cerebellum remapping ratio, and the resulting ratios were averaged across participants of the same dataset to form the test’s statistic. A ratio that significantly deviates from 1 would suggest a difference in relative remapping between the cerebrum and cerebellum. Under the null hypothesis of no difference between cerebral and cerebellar remapping ratios, the cerebral and cerebellar remapping ratios were shuffled within participants and then averaged across participants, a procedure which was repeated 10,000 times to create the null distribution. The position of the true (unshuffled) test statistic within this distribution was then used to obtain a two-sided p-value. Finally, the resulting dataset-specific p-values were tested using Fisher’s method to assess the consistency of effects across the two datasets.

#### Assessing remapping in the putamen

Remapping in the putamen was studied using a more exploratory approach compared to the approach employed for the study of the cerebellar and cerebral cortices. The deprived and intact putamen ROIs were defined based on the Harvard-Oxford probabilistic atlas, at a probabilistic threshold of 90 (see Table 2). Activations evoked by body-part movements in these ROIs were compared between the experimental groups of Dataset2 alone (which had better spatial resolution, allowing the study of this smaller subcortical structure), using two-sided permutation tests, as described for the cerebellar ROI analyses.

#### Confirmatory analysis: spatial layout of body-part representations

Finally, we aimed to confirm previously reported results, by demonstrating differences between the native somatotopy of the cerebellum and the S1/M1 somatotopy (see Introduction). To measure the proximity between native body-part representations, the level of overlap between activations evoked by movements of the hand, lips and feet in the “intact” hemisphere (cerebral hemisphere contralateral to the dominant/intact hand and cerebellar hemisphere ipsilateral to the dominant/intact hand) was measured in all participants using the Dice coefficient (Dice, 1945; Kikkert et al., 2016):

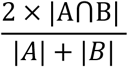

Where A and B represent activations evoked by movements of specific body-parts (intact hand and lips or intact hand and feet) within a sensorimotor mask. To that end, for each participant, the activation maps of intact hand, lips and feet conditions were set to a minimal threshold of at z=2 to allow a relatively wide spread of activation (Kikkert et al., 2016). The few participants who had particularly low spread of activation (<25 voxels; representing 2.5% of the voxels across all analyzed ROIs) in the intact hemisphere in either condition, despite the relatively lenient threshold, were excluded from this particular analysis (Dataset1: 3 control participants; Dataset2: one one-hander and one control participant). In the cerebral cortex, the level of overlapping activations between the hand condition and each of the lips and feet conditions were assessed within a mask of the left pre-central gyrus, taken from the Harvard-Oxford probabilistic atlas (this mask was used without setting a threshold, to contain the central sulcus and both the pre- and post-central gyri, see Figure 8A). In the cerebellar cortex, the level of overlapping activations between the hand condition and each of the lips and feet conditions were assessed within a mask of right lobules I-IV,V and VI, taken from FSL’s cerebellar probabilistic atlas. Each of these three cerebellar masks were thresholded at 50 prior to their unification in order to restrict the unified mask to the sensorimotor sections of the cerebellar anterior lobe (see Figure 8A).

**Figure 3.**
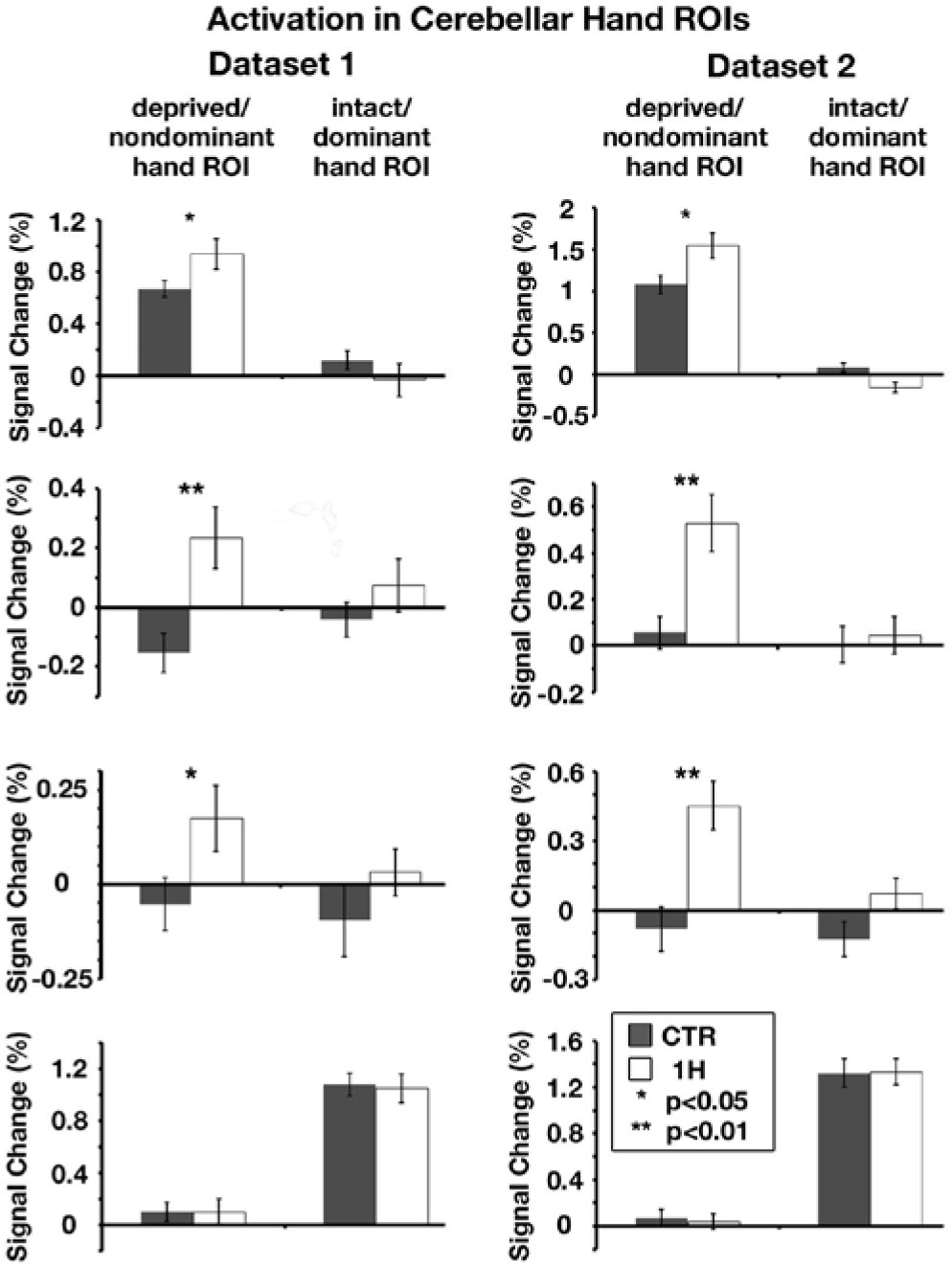
Multiple body-parts activate the deprived cerebellar hand region of one-handers: ROI analysis. The left/right panels show activation levels in Dataset1/Dataset2 (respectively) in the bilateral cerebellar hand regions (independently defined for each dataset, ROIs depicted in Figures 1,2), during residual/nondominant arm (one-handers/controls), lips, feet and intact/dominant hand movements. Activation levels in the deprived cerebellar hand region of one-handers (white bars) were greater than activations in the nondominant hand region of controls (grey bars) in all but the intact hand condition. 1H, one-handers; CTR, controls; intact/dominant hand ROI, ipsilateral to the intact/dominant hand; deprived/nondominant hand ROI, ipsilateral to missing/nondominant hand. Error bars depict SEMs. Results of residual arm movements in a subset of participants from Dataset1 were previously reported (Makin et al., 2013b). The scales of brain activations (y-axes) are not fixed across experimental conditions, to allow better visualization of the inter-group and inter-hemispheric differences within each condition.

**Figure 4.**
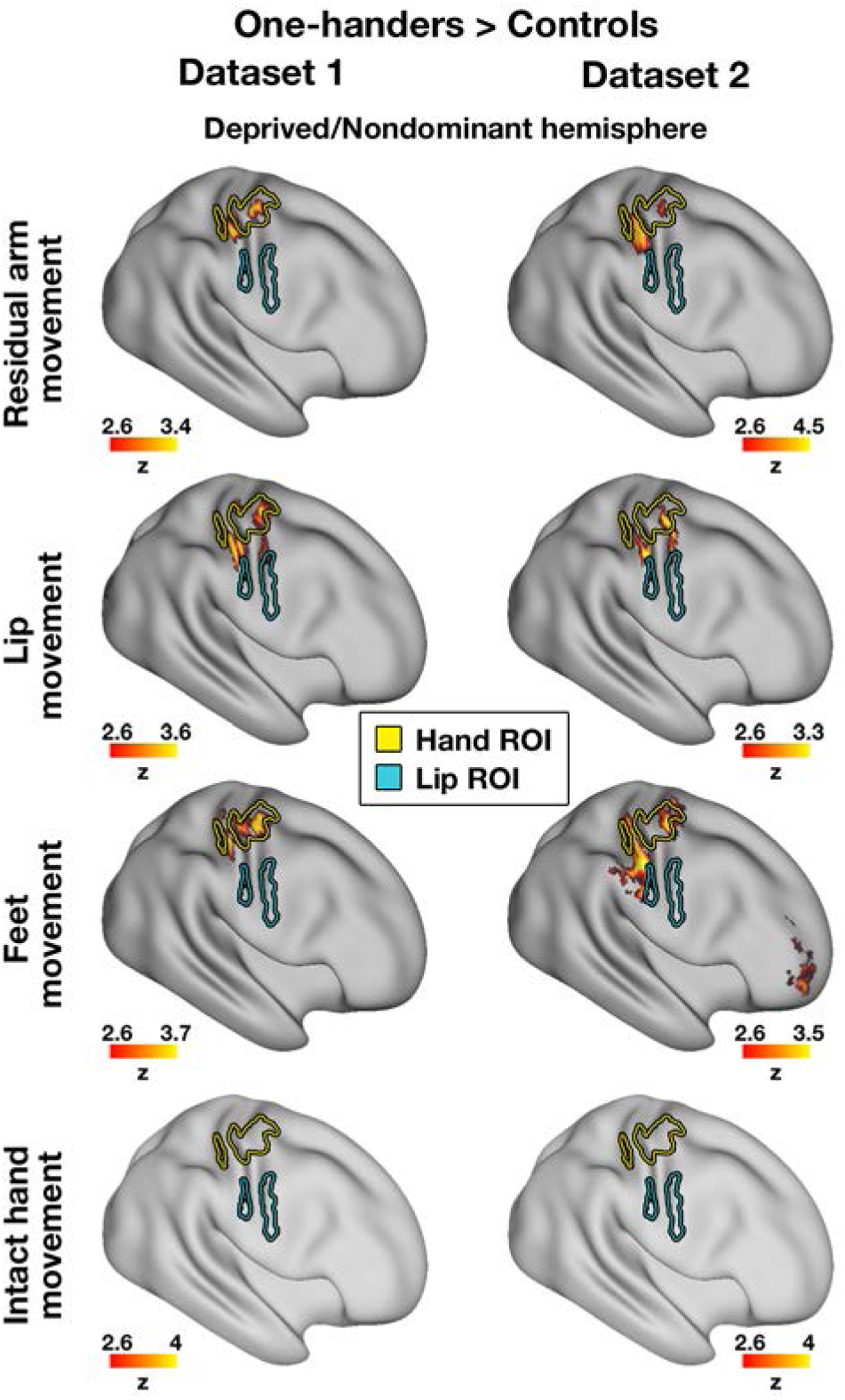
Multiple body-parts activate the deprived cerebral hand-region of one-handers: between-group contrast maps. The left/right panels show between-group contrast maps of Dataset1/Dataset2, respectively, during residual/nondominant arm (one-handers/controls), lips, feet and intact/dominant hand movements, projected onto an inflated surface of a template brain. In each of the arm, lips and feet (but not intact hand) conditions, one-handers showed increased activation compared to controls, centred on the deprived cerebral hand region. Yellow/blue contours indicate the hand/lip ROIs, respectively. Deprived/Nondominant hemisphere, contralateral to missing/nondominant hand. All maps were cluster-based corrected for multiple comparisons across the entire brain. Results of residual arm movements in a subset of participants from Dataset1 were previously reported (Makin et al., 2013b).

**Figure 5.**
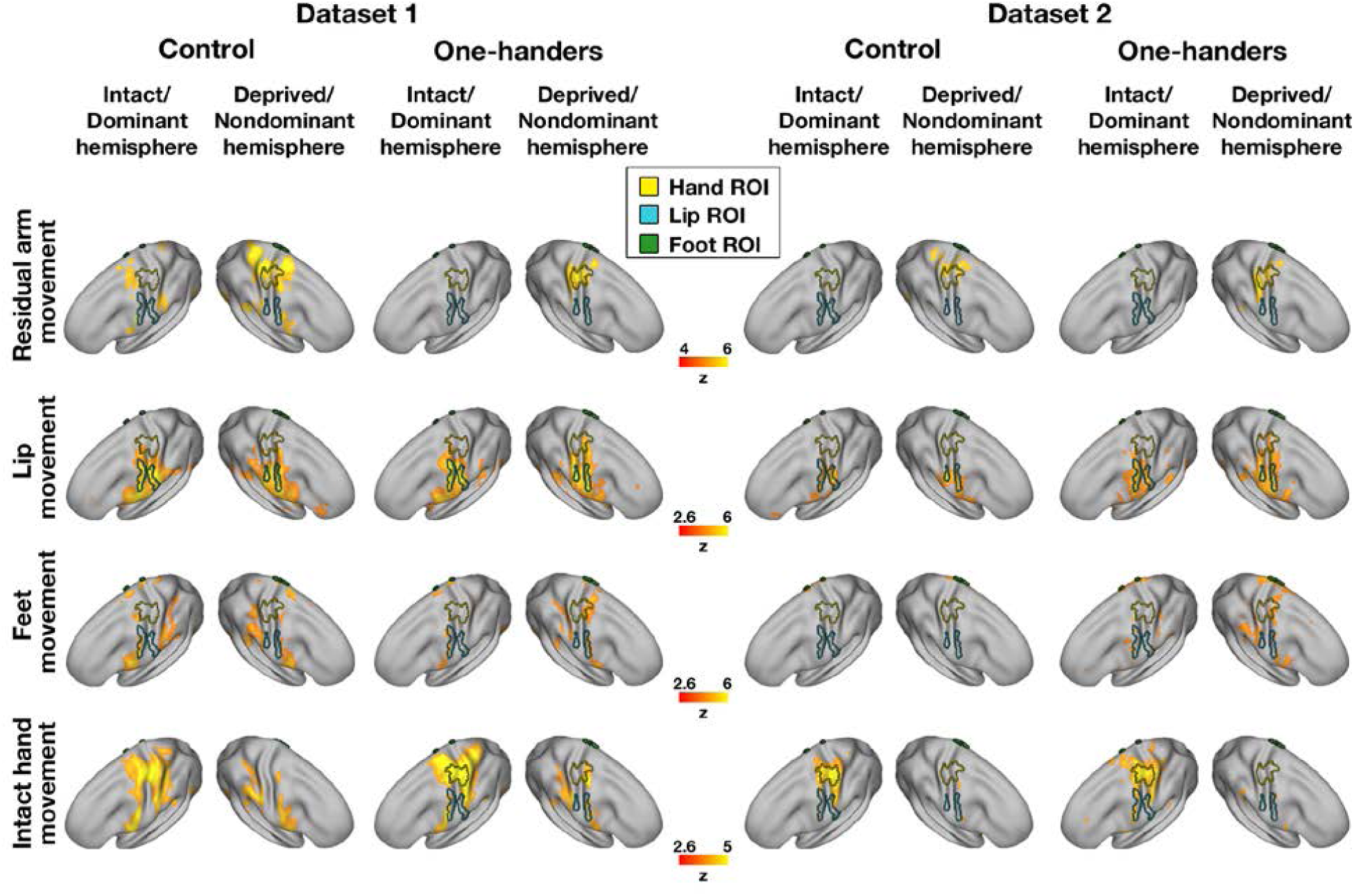
Cerebral within-group activation maps. Within-group activation maps for each experimental condition versus a resting baseline (rows) are presented for the control and one-handed groups of each separate dataset (columns). ROIs were defined based on Dataset3, to ensure full replication of our previously published results in this dataset (Hahamy et al., 2017). All annotations are as in Figure 4. All maps are presented as means of visualisation of the origin of the between-group contrast results presented in Figure 4, and are therefore presented at varying uncorrected thresholds. The within-group maps of one-handers show over-activation in the cerebral deprived hand region in the residual arm, lips and feet conditions, compared to controls. Within-group maps of Dataset3 can be found in Hahamy et al., 2017.

**Figure 6.**
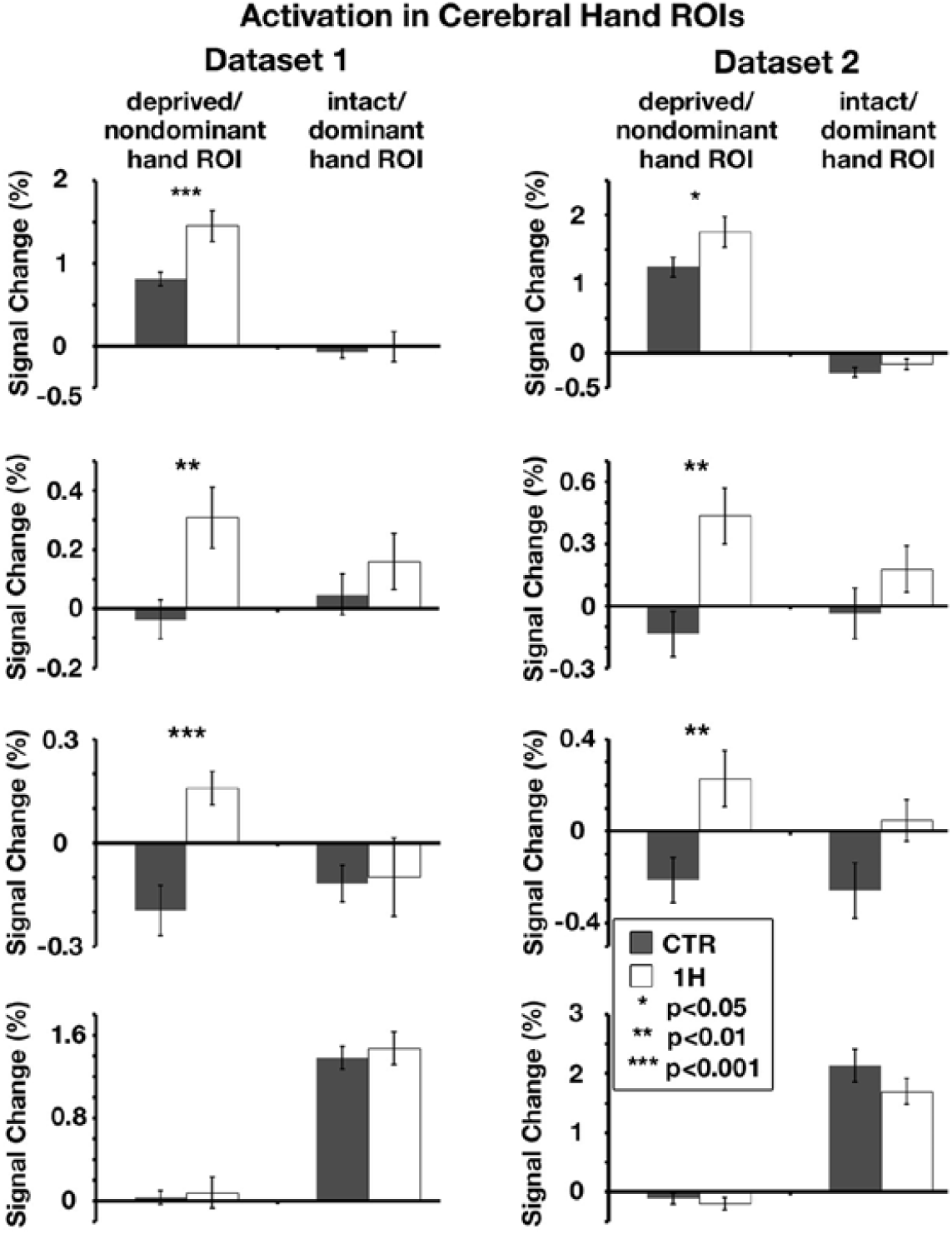
Multiple body-parts activate the deprived cerebral hand-region of one-handers: ROI analysis. The left/right panels show activation levels in Dataset1/Dataset2 (respectively) in the bilateral cerebral hand regions (independently defined, ROIs depicted in Figure 5), during residual/nondominant arm (one-handers/controls), lips, feet and intact/dominant hand movements. Activation levels in the deprived cerebral hand region of one-handers (white bars) were greater than in the nondominant-hand region of controls (grey bars) in all but the intact hand condition. All annotations are as in figure 3. Results of residual arm movements in a subset of participants from Dataset1 were previously reported (Makin et al., 2013b). The scales of brain activations are not fixed across experimental conditions, to allow a better visualization of the inter-group and inter-hemispheric differences within each condition.

**Figure 7.**
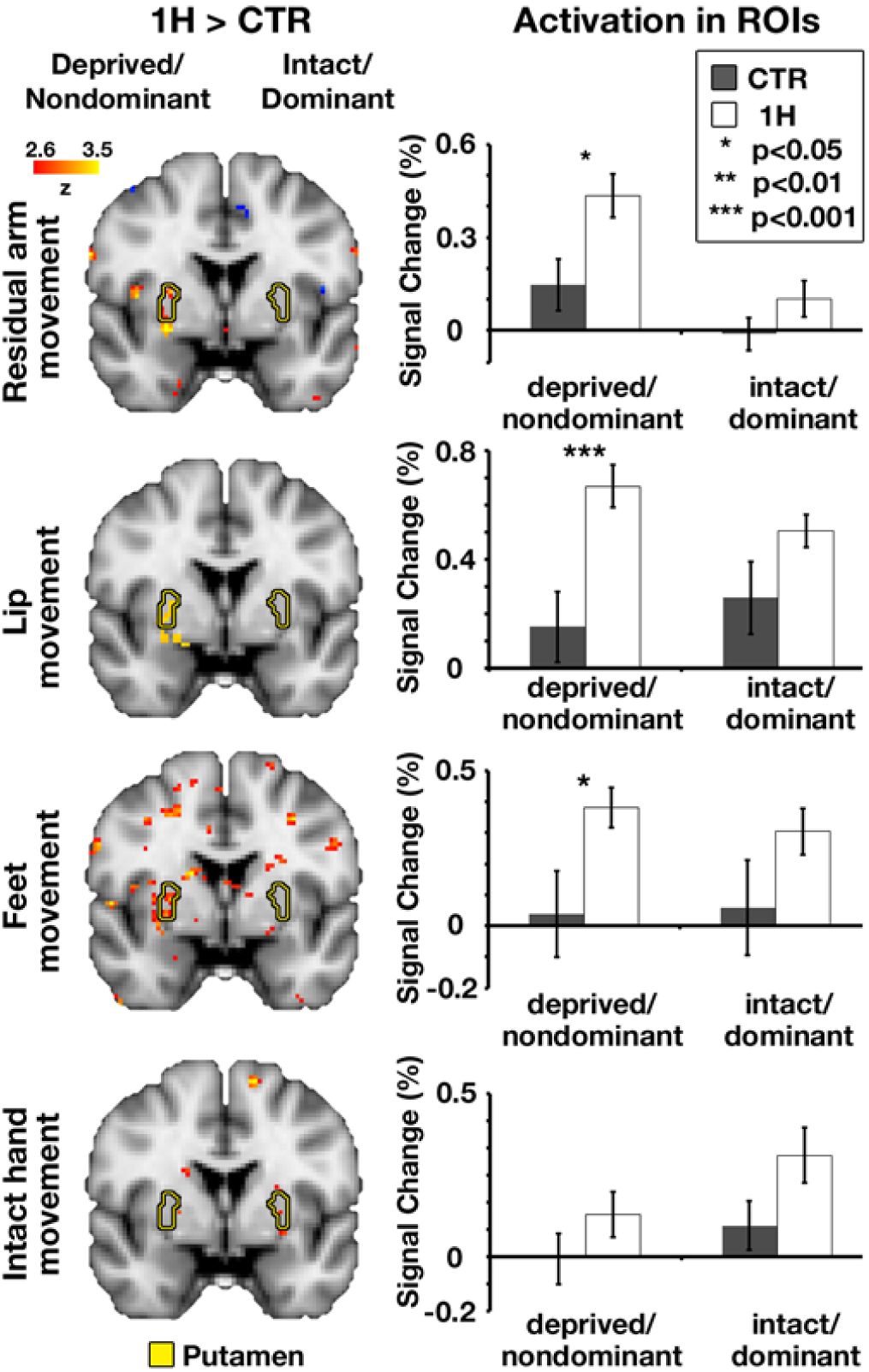
Multiple body-parts over-activated the deprived putamen of one-handers. Left: between-group contrast maps of Dataset2, during residual/nondominant arm (one-handers/controls), lips, feet and intact/dominant hand movements. In each of the arm, lips and feet (but not intact hand) conditions, one-handers showed increased activation compared to controls, centred on the deprived putamen. Yellow contours indicate the bilateral putamen nuclei. Between-group contrast maps of the residual arm, feet and intact hand conditions are presented at an uncorrected threshold of p<0.01. Right: activation levels in Dataset2 in the bilateral putamen nuclei, during each of the experimental conditions. Activation levels in the deprived putamen of one-handers (white bars) were greater than in the nondominant putamen of controls (grey bars) in all but the intact hand condition. All annotations are as in Figure 3.

**Figure 8.**
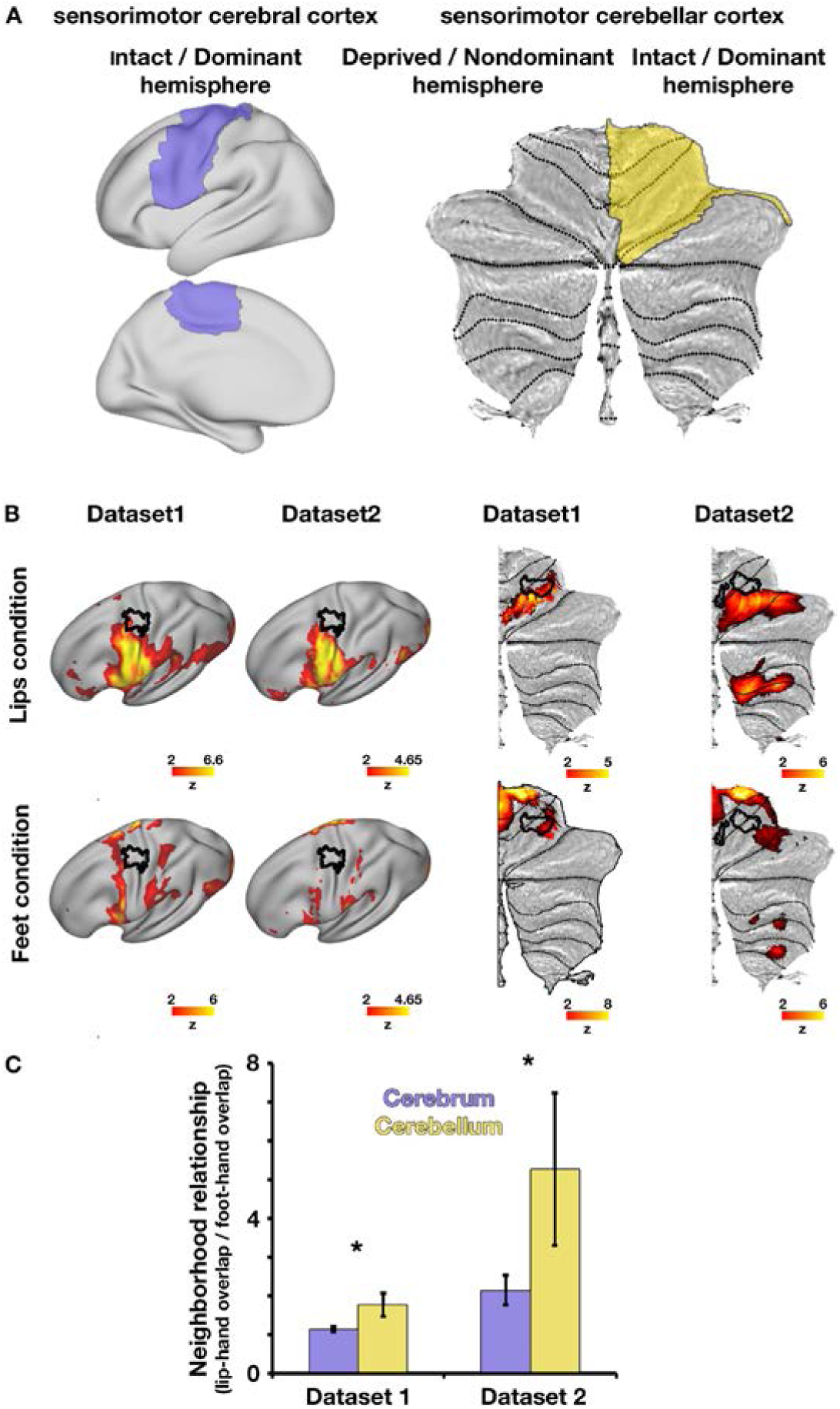
Different overlap relationships of body-part activations between the cerebrum and cerebellum. (A) Sensorimotor masks used to estimate overlap relationships between body-part representations in the intact/dominant cerebral hemisphere (left, marked in purple over an inflated cortical surface) and intact/dominant cerebellar hemisphere (right, marked in yellow over a flattened cerebellar surface). The “intact/dominant” cerebral hemisphere is contralateral to the intact/dominant hand (in controls/one-handers, respectively) and the “intact/dominant” cerebellar hemisphere is ipsilateral to the intact/dominant hand. (B) Averaged maps across all participants of each dataset (columns) in the lips (top row) and feet (bottom row) condition, projected onto surfaces of the intact/dominant cerebral (left) and cerebellar (right) hemispheres. Independent ROIs of the intact/dominant hand are depicted in black contours on these same surfaces. These ROIs are presented for illustration purposes only, and were not used in our statistical analysis of neighborhood relationships, which does not rely on ROIs (see Materials and Methods). (C) Overlap between (i) hand and lip activations and (ii) hand and foot activations were estimated for each participant using the Dice coefficient (see Materials and Methods). The relationship between these overlapping activations was calculated as the ratio between lip-hand overlap and foot-hand overlap (y-axis), which was calculated separately for the cerebrum (purple bars) and cerebellum (yellow bars) within each separate dataset. As evident in both datasets, the ratios of overlap between hand-lip and hand-foot activations are smaller in the cerebellum compared to the cerebral cortex (cerebellar ratios>cerebral ratios), demonstrating different somatotopic layout of body-part representations between the cerebral and cerebellar cortices.

For each participant, 4 Dice coefficients were calculated: overlap between intact hand and feet activations, and overlap between intact hand and lips activations, in each of the cerebrum/cerebellum masks separately. We next aimed to verify that the overlap relationship of body-part representations differs between the cerebrum and the cerebellum, as previously reported (see Introduction). However, a direct comparison between overlap in representations in the cerebrum vs. cerebellum may be confounded by the different spatial scales of these two structures. We therefore targeted a comparison between intra-structure overlap relations, which we will refer to as “neighborhood relationship” of each of the cerebral or cerebellar cortices. This neighborhood relationship was defined as the ratio of lips-hand overlap to feet-hand overlap in each brain structure (cerebrum/cerebellum, Figure 8C). As neighborhood relationships are devised as ratios within each brain structure, they normalize the Dice coefficients and enable a comparison between the cerebrum and cerebellum.

To evaluate whether these neighborhood relationships are different between the cerebrum and cerebellum, a permutation test was employed within each dataset. The test’s statistic was defined as the cross-participants mean ratio between cerebellar neighborhood relationship and cerebral neighborhood relationship (a ratio that significantly deviates from 1 would suggest a difference in topographies between the cerebrum and cerebellum). To this end, for each participant, the cerebellar neighborhood relationship was divided by the cerebral neighborhood relationship. Under the null hypothesis of no difference between cerebral and cerebellar neighborhood relationships, the cerebral and cerebellar neighborhood relationships were shuffled within participants and then averaged across participants, a procedure which was repeated 10,000 times to create the null distribution. The position of the true (unshuffled) test statistic within this distribution was then used to obtain a two-sided p-value. Finally, the resulting dataset-specific p-values were tested using Fisher’s method to assess the consistency of affects across the two datasets.

## Results

### Cerebellar remapping is not restricted by somatotopy

To test whether somatotopy restricts remapping in the cerebellum, we assessed functional remapping of the residual arm (overlapping the hand region, see inserts in Figure1), as well as the lips and feet (whose representations have differing levels of overlap with the hand region, see confirmatory analysis below) in one-handers compared to controls. To this end, we compared results across two independently acquired datasets of one-handers and controls who underwent a functional MRI scan, involving simple movements of the hand, arm, lips and feet. Whole-brain activations evoked by movements of the residual arm, lips and feet (body-part representations previously shown to remap in the cerebral cortex, Hahamy et al., 2017), and of the intact hand (whose representation did not show such remapping, Hahamy et al., 2017) were compared between experimental groups within each dataset. These analyses revealed that movements of the lips and feet, but not movements of the intact hand, excessively activated a region in Lobules V\VI of the cerebellar hemisphere ipsilateral to the missing hand in one-handers, compared to controls (see Figure 1, Table 3). These activation clusters overlapped with an independently defined ROI of the deprived hand region of the anterior cerebellum (see Materials and Methods, Figure 1). Unlike our previous findings in the cerebral cortex, the whole-brain between-group contrast did not reveal increased activation in the cerebellar hand region of one-handers for residual arm movements, compared to controls. This could potentially stem from near complete overlap between the arm and hand representations found in the cerebellum (Mottolese et al., 2013, and see inserts in Figure 1). Specifically, if the arm representation natively overlaps with the hand representation, additional remapping between these representations in one-handers may be too subtle to be detected using a whole-brain analysis. Additional clusters showing increased activation in one-handers compared to controls were also found within specific datasets, but unlike the deprived hand region, these clusters were not consistent across all datasets and task-conditions (see Table 4). The results of the direct between-group-contrasts were further visualized using within-group activation maps of each condition versus a rest baseline (Figure 2).

**Table 3.** Between-group contrast statistics of activation in the hand regions. The number of voxels (#vox), peak intensity (z_max_) and coordinates of the center of gravity of hand-region activations in the cerebellum and cerebrum are presented for each dataset (rows) and task-condition (columns). Coordinates are based on the MNI 152 brain template.

| Brain region | Dataset |  | Residual arm | Lips | Feet |
| --- | --- | --- | --- | --- | --- |
| Cerebellum | 1 | # vox | - | 329 | 246 |
| | | $Z_{\max}$ | - | 5 | 3.94 |
|  |  | Coordinates | - | -13,-55,-17 | -21,-58,-16 |
|  | 2 | # vox | - | 594 | 623 |
| | | $Z_{\max}$ | - | 4.45 | 4.03 |
|  |  | Coordinates | - | -17,-59,-21 | -21,-57,-22 |
| Cerebrum | 1 | # vox | 244 | 439 | 268 |
| | | $Z_{\max}$ | 3.82 | 4.5 | 4.11 |
|  |  | Coordinates | 46,-20,58 | 45,-17,54 | 44,-20,58 |
|  | 2 | # vox | 422 | 411 | 450 |
| | | $Z_{\max}$ | 5.79 | 4.24 | 4.14 |
|  |  | coordinates | 49,-20,56 | 48,-16,54 | 52,-22,50 |

**Table 4.** Between-group contrast statistics of increased activation in one-handers compared to controls outside the hand regions. The number of voxels, peak intensity (z_max_) and coordinates of the center of gravity of significant activation clusters are presented for each dataset and task-condition. Coordinates are based on the MNI 152 brain template.

| **Dataset** | **condition** | **Region** | **Region** | **Region** |
| --- | --- | --- | --- | --- |
| **1** | Lips | supracalcarine cortex,<br>735 voxels,<br>$Z_{\max}=3.75$,<br>(2,-79,13) | Deprived \<br>nondominant occipital<br>pole,<br>342 voxels,<br>$Z_{\max}=4.01$,<br>(20,-91,23) | Intact \ dominant<br>parahippocampal gyrus,<br>217 voxels,<br>$Z_{\max}=3.9$,<br>(-26,-10,-23) |
| **2** | Lips | Deprived \ nondominant<br>putamen,<br>288 voxels,<br>$Z_{\max}=3.7$,<br>(26,-4, -1) | Intact \ dominant<br>parahippocampal gyrus,<br>187 voxels,<br>$Z_{\max}=3.83$,<br>(-30,-17,-28) | Deprived \ nondominant<br>cerebellar lobule VIIb,<br>167 voxels,<br>$Z_{\max}=4$,<br>(-7, -78, -48) |
| | Feet | Intact \ dominant frontal<br>pole,<br>221 voxels,<br>$Z_{\max}=4.18$,<br>(-27,57,0) | Deprived \<br>nondominant frontal<br>pole,<br>177 voxels,<br>$Z_{\max}=3.65$,<br>(29,49,7) | |

We next aimed to measure the degree of remapped activation in the deprived cerebellar hand region during movements of different body parts, and assess its consistency across datasets. To this end, within each dataset and movement condition separately, between-group permutation tests were used to compare mean fMRI activation values (percent signal change) obtained from two independent hand ROIs (deprived and intact hand regions of the cerebellar hemispheres’ anterior lobe, ROIs depicted in Figures 1,2). These ROIs were obtained from the control group of one dataset and tested on the other dataset, and are completely independent of the between-group contrast analysis reported above. Results of these tests were combined across the two datasets (See Materials and Methods; Dataset-specific p-values for all experimental conditions are presented in Table 5). As shown in Figure 3, these analyses confirmed increased activation in the deprived cerebellar hand ROI when one-handers moved their lips (χ^2^_(4)_ =23.21, p<0.001, α=0.017) and feet (χ^2^_(4)_ =19.91, p<0.001, α=0.017), as well as their residual arm (χ^2^_(4)_ =15.29, p=0.004, α=0.017 Bonferroni corrected, Fisher’s method for all tests), compared with controls. Movements of the intact hand (whose representation does not remap in the cerebral cortex, Hahamy et al., 2017) did not result in increased activation in the deprived cerebellar hand region of one-handers (χ^2^_(4)_ =3.33, p=0.51, Fisher’s method). In addition, two-way interactions were consistently revealed between hemispheres and groups (non-dominant/residual arm: χ^2^_(4)_ =20.61, p<0.001; lips: χ^2^_(4)_ =19.39, p<0.001; feet: χ^2^_(4)_ =19.23, p<0.001, Fisher’s method, α=0.017 Bonferroni corrected for all tests). These interactions reflect dissociated recruitment of the deprived cerebellar hand region by movements of various body parts in one-handers, in comparison with the intact cerebellar hand region and with the control group (Figure 3). These findings echo the pattern of remapping we previously reported in the cerebral cortex of one-handers, and reflect sensorimotor remapping which is not limited to the immediate neighbors overlapping with the deprived hand region.

**Table 5.**
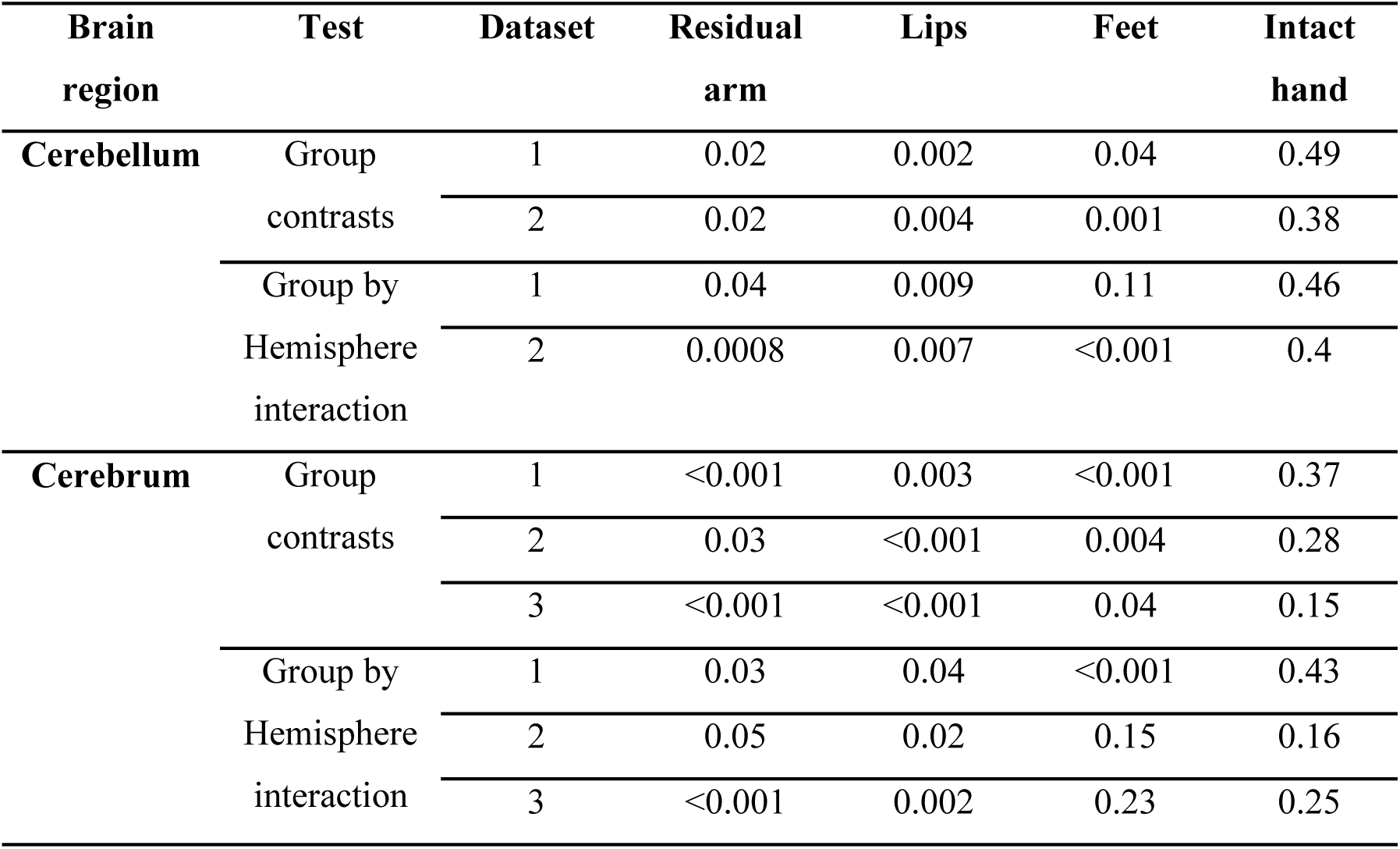
Dataset-specific p-values per brain region and experimental condition. Dataset-specific p-values (rows) are derived from permutation tests for each experimental condition (columns) for both cerebellar and cerebral hand ROIs (top\bottom of table, respectively). Results of Dataset3 were previously reported in Hahamy et al. 2017. Results for the residual arm condition in a subsample of participants from Dataset1 were reported in Makin et al. 2013.

To further evaluate the interplay between remapping and somatotopy, the remapping levels of the lip and foot representations in one-handers were directly compared. If somatotopy drives remapping, the lip representation, which overlaps with the deprived hand representation, should show more remapping compared to the foot representation, which does not overlap with the hand region (see also Confirmatory analysis below). However, despite different levels of overlap with the cerebellar hand region, no difference was found between lip and foot remapping into this region (Dataset1: p=0.33, Dataset2: p=0.32, permutation tests). Furthermore, a 3-way ANOVA with factors group (controls/one-handers), hemisphere (intact/deprived) and body-part (lips/feet) revealed no 3-way interaction (Dataset1: F(1,36)=1.86, p=0.18, Dataset2: F(1,24)=0.47, p=0.5). This indicates that somatotopic proximity does not determine the degree of remapping. Note, however, that these null results do not allow a formal interpretation.

### Similar pattern of remapping seen in the cerebellar and cerebral deprived hand regions of one-handers

We next used these two datasets to test the reproducibility of our previous findings of remapping in the cerebral cortex of one-handers (our previously reported results were based on Dataset3, and can be found in Hahamy et al., 2017). As shown in Figure 4 (and further visualized in Figure 5), movements of the residual arm, lips or feet, but not movements of the intact hand, activated the deprived cerebral S1/M1 hand region to a greater extent in one-handers compared to controls, as shown using whole-brain between-group contrast maps (also see Table 3). Additional clusters showing increased activation in one-handers compared to controls were also found within specific datasets, but unlike the deprived hand region, these clusters were not consistent across all datasets and task-conditions (see Table 4). These results were further supported by ROI analyses (ROIs presented in Figures 4,5). To assess reproducibility, the results of the ROI analyses were combined across the two current datasets as well as the dataset used in our previous study (dataset-specific p-values for all experimental conditions are presented in Table 5). As depicted in Figure 6 and in Table 6, these tests confirmed increased activation in the deprived cerebral hand ROI when one-handers moved their residual arm, lips and feet compared with controls. Group by hemisphere interactions consistently revealed dissociated recruitment of the deprived cerebral hand region (compared to the intact hand region) by movements of various body parts between one-handers and controls (Table 6). Table 7 presents the results of integration across datasets using additional meta-analysis measures for both the cerebral and cerebellar hand regions.

**Table 6.**
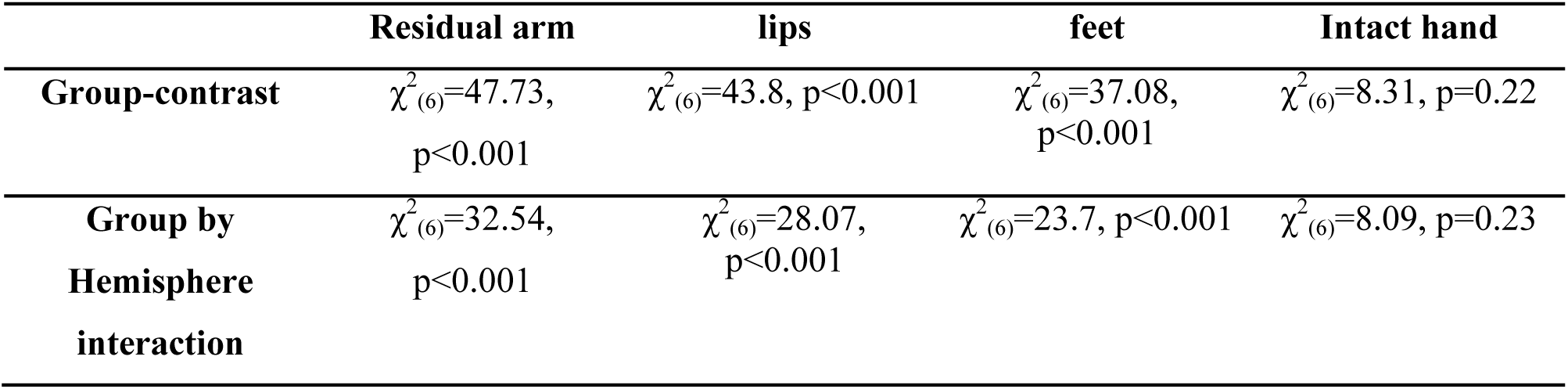
Meta-analysis statistics for cerebral cortex activations. Fisher’s method statistics (χ2 and p-value) are presented for the between-group contrasts and group by hemisphere interactions (rows) across experimental conditions (columns). All p-values are Bonferroni corrected, α=0.017.

|  | **Residual arm** | **lips** | **feet** | **Intact hand** |
| --- | --- | --- | --- | --- |
| **Group-contrast** | $\chi^2_{(6)}=47.73$,<br>p<0.001 | $\chi^2_{(6)}=43.8$, p<0.001 | $\chi^2_{(6)}=37.08$,<br>p<0.001 | $\chi^2_{(6)}=8.31$, p=0.22 |
| **Group by Hemisphere interaction** | $\chi^2_{(6)}=32.54$,<br>p<0.001 | $\chi^2_{(6)}=28.07$,<br>p<0.001 | $\chi^2_{(6)}=23.7$, p<0.001 | $\chi^2_{(6)}=8.09$, p=0.23 |

**Table 7.**
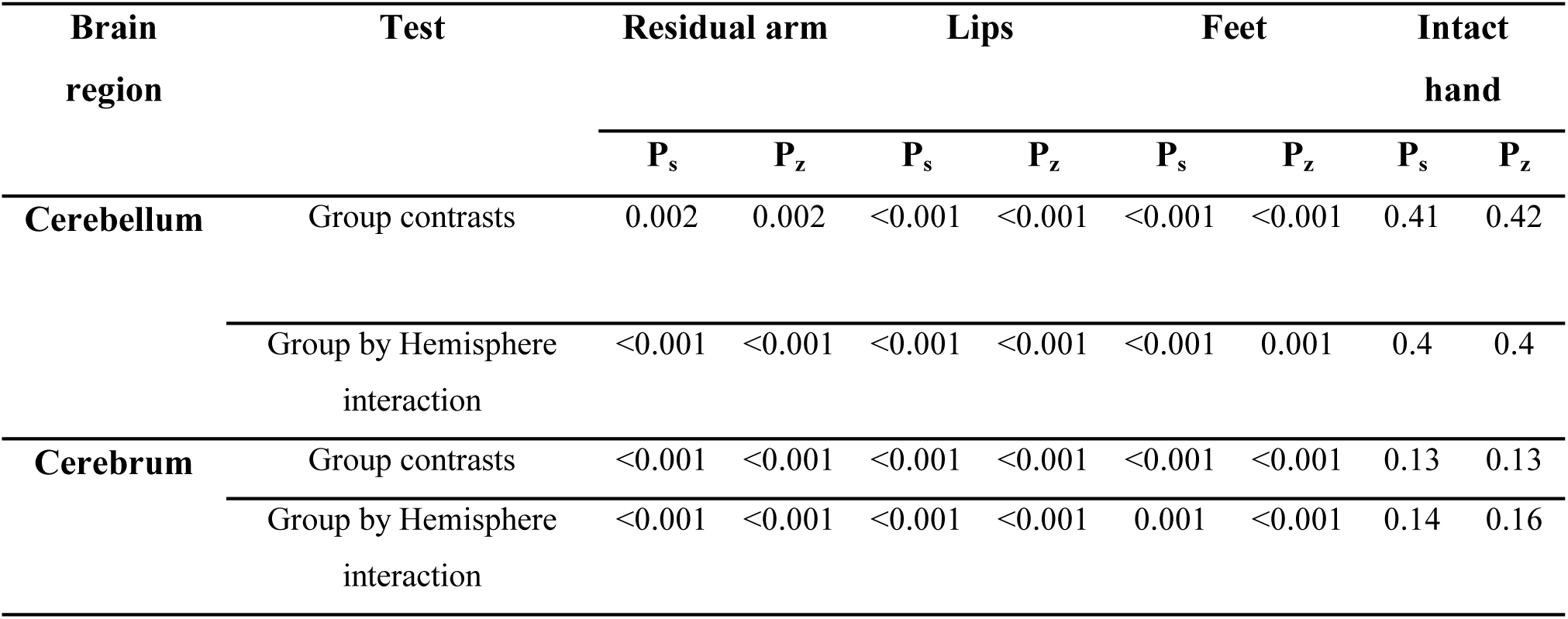
Assessment of the consistency of results across datasets using additional meta-analysis methods. Results are based on the Stouffer’s test (Stouffer et al., 1949) and the weighted Z-test (weights set to the square root of each sample size, Liptak, 1958). Resulting p-values (P_s_ denotes Stouffer’s test and P_z_ denotes the weighted Z-test, α=0.0125 Bonferroni corrected) are presented for the cerebrum and cerebellum hand regions (rows) and for each experimental condition (columns).

| Brain region | Test | Residual arm |  | Lips |  | Feet |  | Intact hand |  |
| --- | --- | --- | --- | --- | --- | --- | --- | --- | --- |
| | | $P_s$ | $P_z$ | $P_s$ | $P_z$ | $P_s$ | $P_z$ | $P_s$ | $P_z$ |
| Cerebellum | Group contrasts | 0.002 | 0.002 | <0.001 | <0.001 | <0.001 | <0.001 | 0.41 | 0.42 |
|  | Group by Hemisphere interaction | <0.001 | <0.001 | <0.001 | <0.001 | <0.001 | 0.001 | 0.4 | 0.4 |
| Cerebrum | Group contrasts | <0.001 | <0.001 | <0.001 | <0.001 | <0.001 | <0.001 | 0.13 | 0.13 |
|  | Group by Hemisphere interaction | <0.001 | <0.001 | <0.001 | <0.001 | 0.001 | <0.001 | 0.14 | 0.16 |

To evaluate whether different somatotopic layouts would relate to different patterns of body-part remapping, we compared the remapping seen in the deprived cerebellar and deprived cerebral hand regions of one-handers (see Material and Methods). This analysis revealed that, although the cerebellum and cerebrum have different somatotopic layouts (see also Confirmatory analysis below), the lip to foot remapping ratios did not differ between these two sensorimotor terminals (Dataset1: p=0.09, Dataset2: p=0.15, permutation tests; χ^2^_(4)_ =8.6, p=0.072, Fisher’s Method). But as noted above, null results should be interpreted with caution.

### Remapping in the Putamen

Since the putamen has previously been shown to contain a somatotopic map (Nambu, 2011), and since the basal ganglia has reciprocal connections with both the cerebellum and the primary sensorimotor cortex (Nambu, 2011; Dum et al., 2014; Zeharia et al., 2015), we wished to explore remapping patterns in this terminal. A challenge in studying this area is that its somatotopy is substantially more compact than that of the cerebellum, requiring increased spatial resolution, which was only available in Dataset2. The left panel of Figure 7 depicts uncorrected between-group-contrast maps of Dataset2. Similar to our results in the cerebellum and S1/M1, these maps demonstrated increased activation in the deprived putamen in the residual arm, lips and feet conditions, but not in the intact hand condition in one-handers compared to controls. Only results of the lips condition survived correction for multiple comparisons over the whole brain (Table 4).

ROI analysis provides a more sensitive test for remapping. As depicted in the right panel of Figure 7, these tests revealed significantly increased activation in the deprived putamen in the residual arm (p=0.01), lips (p=<0.001) and feet conditions (p=0.02; permutation tests, α=0.05), but not in the intact hand condition (p=0.1). These effects were accompanied by near-significant group (controls/one-handers) by hemisphere (intact/deprived putamen) interactions (residual arm: p=0.048, lip: p=0.007, feet: p=0.057, permutation tests, α=0.05), demonstrating the specificity of this effect to the deprived putamen. Note that, unlike in the cerebrum and cerebellum, activation in the putamen tends to be bilateral (Gerardin et al., 2003), hence the non-negligible activation levels in both hemispheres.

### Confirmatory analysis: neighborhood relationship between body-part representations differ between the cerebrum and cerebellum

In both the cerebral and cerebellar cortices, the hand region resides between the foot and lip regions, however, the level of overlap between these representations was previously reported to differ between the two brain structures (see Introduction). To confirm this difference in overlap between body-part representations (hereafter, neighborhood relationship) in the cerebrum and cerebellum, we studied the overlap in activations evoked by movements of these body-part in controls and one-handers’ intact hemisphere (notice that no between-group differences were found in the intact hemisphere, Figures 1&4).

We confined activations to the sensorimotor parts of the cerebrum and cerebellar anterior lobe (Figure 8A), and employed the Dice coefficients (Dice, 1945; Kikkert et al., 2016; see Materials and Methods). As demonstrated in the intact/dominant hemispheres of controls and one-handers in Figure 8B, some degree of overlap was indeed observed between the peripheral aspects of the hand region and the activations evoked by lips and feet movements in both cerebrum and cerebellum. This level of overlap was evaluated using permutation tests on each of the dataset-specific Dice coefficients. Results of these tests were then combined across the two datasets (See Materials and Methods). This analysis demonstrated differences in neighborhood relationships between the cerebrum and cerebellum (Dataset1 p=0.02, Dataset2 p=0.048 permutation tests; Meta-analysis: χ^2^_(4)_ =13.59, p=0.009 Fisher’s method, α=0.05), reflecting that the representations of the lip and foot show more similar levels of overlap with the hand representation in the cerebral cortex (Makin et al., 2015) relative to the cerebellum (Figure 8C).

## Discussion

Here we report large scale remapping of body-part representations in both the cerebellar and cerebral cortices of individuals born without one hand, and provide similar preliminary results in the putamen. In all terminals, the residual arm, lips and feet activated the deprived hand region (Figures 1, 4 & 7). Remapping was specific to the missing hand regions of these terminals (as reflected in our whole-brain analyses and significant group by hemisphere interactions), despite differences in the somatotopic layouts across these sensorimotor terminals (Manni and Petrosini, 2004; Mottolese et al., 2013; Makin et al., 2015; Mottolese et al., 2015). Our findings therefore challenge the view that sensorimotor remapping is restricted by the underlying somatotopy of the remapped regions (Merzenich et al., 1984; Pons et al., 1991; Merzenich and Jenkins, 1993; Faggin et al., 1997; Florence et al., 1998; Margolis et al., 2012; Striem-Amit et al., 2018), at least following congenital deprivation.

Previous studies of similar sensorimotor-deprived populations, relying on relatively small sample sizes, produced mixed evidence for the extent and drivers of remapping. Here we used a large imaging database of one-handers (n=26), and demonstrated the reproducibility of our main results across independently-acquired datasets, thereby establishing statistical validity (Ioannidis et al., 2014; Picciotto, 2018). Our findings therefore contribute robust evidence that remapping extends beyond the boundaries of the somatotopy, and emphasize the need to consider sensorimotor remapping following congenital malformation as a more complex phenomenon than has previously been discussed. Future large-scale studies of both functional representation and connectivity, as well as stimulation studies (e.g. transcranial magnetic stimulation, Stoeckel et al., 2009) will be needed to fully understand the functional specificity and underlying factors that derive the reported remapping.

If remapping into the deprived hand region is not exclusively restricted to the neighboring representations, what other factors determine which representation undergoes remapping and which does not? One possibility is that remapping is shaped by altered inputs to the deprived cortex, due to compensatory behavior. We have previously characterized the behavioral repertoire of one-handers, which comprises of utilization of their residual arm, lips and feet to compensate for their hand absence (Hahamy et al., 2017). As previously reported and further extended here, the same body-parts used for compensatory purposes also remap onto the deprived hand regions of both the cerebellum and cerebrum. Furthermore, the intact hand, which is not over-used for compensatory purposes in one-handers (Makin et al., 2013b; Philip and Frey, 2014; Hahamy et al., 2017), does not show remapping onto either the cerebellar or cerebral deprived hand regions. However, we cannot exclude the possibility that remapping in the deprived hand region of one-handers is restricted to body-part representations within the deprived hemisphere. This is because in the current experimental design, the intact hand is the only body-part contralateral to the deprived cerebral hemisphere/ipsilateral to the deprived cerebellar hemisphere.

It is important to mention that so far we have been unable to identify a correlation between individuals’ idiosyncratic compensatory strategies and brain remapping. Moreover, other studies reported large-scale remapping dissociated from compensatory behavior in congenital or juvenile bilateral hand loss (Yu et al., 2014; Striem-Amit et al., 2018). For example, Striem-Amit and colleagues (2018) recently demonstrated that body-part representations neighboring the deprived hand region can show remapping, even if these body-parts are not prominently used for compensatory purposes. These discrepancies across studies could be attributed to the fact that compensatory daily behavior is difficult to quantify comprehensively and reliably. Alternatively, it could be speculated that the development of one-handers’ intact hand grants the missing hand region some sensorimotor scaffolding relating to hand functionality (e.g. via inter-hemispheric functional connectivity, Hahamy et al., 2017). This, in turn, may guide remapping in one-handers based on behavioral criteria (e.g. relevance for supporting the intact hand), which will not be available or functionally relevant following bilateral hand malformation. We also cannot exclude the possibility that behavior and brain remapping may not be directly related. For example, the observed remapping in one-handers may merely reflect weak normal inputs from different body-parts to the hand region, which are typically supressed. In the absence of a hand, these inputs may simply be unmasked, and not necessarily causally support compensatory behavior (for further discussion see Krakauer and Carmichael, 2017; Makin and Bensmaia, 2017). Taken together, further research is needed to validate the causal origins and consequences of behavior on the large scale remapping reported here.

Similar controversy regarding the role of somatotopic boundaries in shaping remapping also exists in amputation research. In adult amputees, remapping is commonly attributed to residual arm representation (Kew et al., 1994; Irlbacher et al., 2002; Raffin et al., 2016); (but see Gagne et al., 2011; Makin et al., 2013b) and mouth representation (Flor et al., 1995; Elbert et al., 1997; Karl et al., 2001; Lotze et al., 2001; MacIver et al., 2008; Foell et al., 2014); (though see Makin et al., 2013a; Makin et al., 2015; Raffin et al., 2016), both thought to neighbor the hand region (though see Zeharia et al., 2015; Roux et al., 2018; as well as hand and lip regions in Figure 4). More recent findings reveal remapping of the intact hand representation into the deprived cortex (Bogdanov et al., 2012; Makin et al., 2013b; Philip and Frey, 2014). These more recent findings have ascribed remapping to the compensatory use of amputees’ intact hand. Thus, findings across varied sensorimotor-deprived populations raise the possibility that body-part representations that have little overlap, if any, with the hand region (lips and feet in one-handers, feet in individuals with bilateral congenital limb loss or childhood amputation, and intact hand in amputated adults) can remap into the deprived hand region.

Although we discuss commonalities in remapping across the life-span, this is not to imply that remapping bears the same mechanistic and functional meaning when occurring at different life stages. Hand function begins to form *in utero* (Zoia et al., 2007) and continues to develop into late childhood (Kuhtz-Buschbeck et al., 1998). As such, congenital hand malformation offers multiple opportunities for functional remapping during development. Indeed, vast research on visual and auditory deprivations introduced the notion of the critical period - an early period of life in which sensory experience may have greater impact on brain remapping and consequent behavior, compared to later periods (for review, see Kral, 2013; Voss, 2013). In contrast, classical research of sensorimotor deprivation documented extensive remapping in adults (Pons et al., 1991; Florence et al., 1998). Although the extent and functional significance of remapping in later life are still debated (Collignon et al., 2013; Bedny, 2017; Makin and Bensmaia, 2017; Singh et al., 2018), it is worth noting that in comparison to congenital blindness and deafness research, sensorimotor deprivation is confined to the sensorimotor network, and is thus smaller in scale. Therefore, amputation-related deprivation might provide more opportunities/restrictions for remapping across the life-span, meaning sensorimotor remapping may still be feasible in adulthood (Dempsey-Jones et al., 2019).

Finally, the remapping reported here may indeed be constrained by proximity between body-part representations - not in the cerebellum/cerebrum, but rather in subcortical sensorimotor terminals. While it has originally been suggested that sensorimotor remapping occurs at the level of the cerebral cortex (Pons et al., 1991; Florence et al., 1998), recent studies in monkeys emphasize the role of subcortical structures, such as the brainstem, in which the layout of somatotopic representations differs from that of the cerebral cortex (Jain et al., 2000; Kambi et al., 2014; Chand and Jain, 2015; Liao et al., 2016). For example, Kambi and colleagues (2014) demonstrated that facial remapping in the deprived cerebral hand region of spinal-cord-injured monkeys is abolished upon inactivation of the deprived cuneate nucleus. The fact that the cuneate nucleus does not normally receive inputs from the face suggests that remapping seen in the cerebral cortex is likely driven by reorganisation at the level of the brainstem (see also Herbert et al., 2015 for a related example of remapping in the motor cortex). It is therefore plausible that the remapping we observed in one-handers may also be initiated in upstream sensorimotor terminal.

Indeed, our data provide initial evidence for remapping in one-handers’ putamen, which mirrors the remapping patterns of the cerebral and cerebellar cortices. Interestingly, representations of the hand, lip and foot, which are distant in the cerebral/cerebellar cortices, neighbor each other in the putamen of the human basal ganglia (Gerardin et al., 2003; Staempfli et al., 2008). These neighboring representations may thus more easily remap onto the deprived hand region, and consequently, projections from the putamen hand representation to its cerebral/cerebellar counterparts would appear as remapping that is independent of the cerebral/cerebellar somatotopic layouts. This hypothesis is consistent with anatomical evidence for a closed-loop, reciprocal circuit between primary sensorimotor cortex, cerebellum and basal ganglia, such that each terminal projects to and receives inputs from each other terminal, with varying somatotopic layouts within each terminal (Nambu, 2011; Dum et al., 2014; Zeharia et al., 2015). As our findings reveal remapping in all three of these interconnected terminals, it is plausible that the documented remapping is initiated by the basal ganglia, or further upstream. However, the results we reported in the putamen could only be observed in one dataset, and lacked the spatial resolution to accurately allocate the position of the putamen hand region. These results therefore await further confirmation with more specialised data collection tools. In addition, since our results demonstrate remapping in both primary somatosensory and motor cortices (Figure 4), which have differing upstream hierarchies, the role of subcortical structures in driving cortical remapping requires further research. Nevertheless, it is tempting to speculate that the upstream sensorimotor terminal at which remapping may be initiated would contain a somatotopic layout specifically suitable for supporting the emerging repertoire of compensatory behaviors.

## Acknowledgements

We thank Alona Cramer, Jan Scholtz, Fiona van der Heiligenberg, Harriet Dempsey Jones and Lucilla Cardinali for help in data collection, and Victoria Root and Arabella Bouzigues for comments on the manuscript. We thank Opcare for help with participants recruitment, and our participants for taking part in these studies.

